# Impact of increased membrane realism on conformational sampling of proteins

**DOI:** 10.1101/2021.03.23.436674

**Authors:** Austin T. Weigle, Matthew Carr, Diwakar Shukla

## Abstract

The realism and accuracy of lipid bilayer simulations through molecular dynamics (MD) is heavily dependent on the lipid composition. While the field is pushing towards implementing more heterogeneous and realistic membrane compositions, a lack of high-resolution lipidomic data prevents some membrane protein systems from being modeled with the highest level of realism. Given the additional diversity of real-world cellular membranes and protein-lipid interactions, it is still not fully understood how altering membrane complexity affects modeled membrane protein function or if it matters over long timescale simulations. This is especially true for organisms whose membrane environments have little to no computational study, such as the plant plasma membrane. Tackling these issues in tandem, a generalized, realistic, and asymmetric plant plasma with more than 10 different lipid species membrane is constructed herein. Classical MD simulations of pure membrane constructs were performed to evaluate how altering the compositional complexity of the membrane impacted the plant membrane properties. The *apo* form of a plant sugar transporter, OsSWEET2b, was inserted into membrane models where lipid diversity was calculated in either a size-dependent or -independent manner. An adaptive sampling simulation regime validated by Markov-state models was performed to capture the gating dynamics of OsSWEET2b in each of these membrane constructs. In comparison to previous OsSWEET2b simulations performed in a pure POPC bilayer, we confirm that simulations performed within a native-like membrane composition alter the stabilization of *apo* OsSWEET2b conformational states by ~1 kcal/mol. The free energy barriers of intermediate conformational states decrease when realistic membrane complexity is simplified, albeit roughly within sampling error, suggesting that protein-specific responses to membranes differ due to altered packing caused by compositional fluctuations. This work serves as a case study where a more realistic bilayer composition makes unbiased conformational sampling easier to achieve than with simplified bilayers.

## INTRODUCTION

Biological membranes of natural origin are nonhomogeneous, multicomponent and asymmetric with a large diversity of lipid chemical structures. Resulting protein-lipid interactions are as diverse as the membrane bilayer composition itself and deserve further study. Although recent realistic membrane studies have emerged, the modeling of pure complex membranes and their interactions with membrane proteins is still in a characterization phase. Molecular dynamics (MD) simulations account for the positions of all atoms of a given system at any moment in time, providing an exceptional level of atomistic, label-free resolution for studying membrane properties and membrane protein function. As the popularity of MD has grown in parallel to the advent of more efficient computing capabilities, modern practices concerning the simulation of membrane systems have also advanced.^1,2^ Specifically, increasing attention has been paid to appropriate modeling of membrane realism in the hopes of better capitulating native biological processes.^3^ How “realistic” a membrane’s simulated behavior can be is influenced by the selection of forcefields, system setup tools, and the constituent lipid diversity representing the modeled cellular environment. System preparation protocols and forcefield development behind the assembly and parameterization of membrane-containing MD systems have continuously improved over the last decade.^4,5^ The refinement of these necessary components of membrane system setup have resulted in increased predictive power; for instance, coarse-grained models are widely accepted for accurately predicting lipid-protein interactions.^3,6,7^ Still, uncertainty surrounds how realistic any membrane composition should be for a given MD application, where the benefit of “better” lipid representation is not fully understood.

Simulation of realistic membranes is not yet a standard modeling convention for a number of reasons. Firstly, high-resolution lipidomic data is not available for all cell types of all organisms to reveal accurate chemical diversity nor leaflet asymmetry. This is mainly due to a lack of interest in resolving the complete “lipidomic picture” for a diversity of organisms’ native membrane environments or because spectrometry technology was previously not advanced enough to fully resolve the identity of both lipid acyl chains when first studied. Given crystal structure availability of biodiverse origins without corresponding membrane characterizations available, MD practitioners resort to incorporating simplified membranes into their models. Up until the past decade, this practice of using simplified models has resulted in regularly unchallenged “default” modeling practices. For instance, single component mammalian membranes are commonly represented as POPC, and bacterial membranes are represented by POPE/POPG. Meanwhile, realistic model bilayers include at least six different lipid species.^3^

Secondly, the modeling of an asymmetric membrane is computationally difficult, due to both leaflets being constrained by the same box volume and pressure coupling. Compounded by a lack of data validating explicit leaflet-specific ratios of lipid species, determining how different lipid types should be allotted between the upper and lower leaflets often results from tedious trial and error. Some methods have been developed to make the construction of asymmetric membranes become more systematic,^8–12^ but these approaches do not completely eradicate the lateral pressure differences inherent to leaflet asymmetry.^13^ Additionally, the majority of realistic membranes vary the number of different lipid species between two to six different lipids,^3^ minus a few exceptions.^14–19^ Again, the question of how complex a realistic membrane should be remains unanswered. Taken together, the inherent challenges and increased computational costs involved in validating realistic membrane modeling explain why relatively few computational studies have incorporated such complex bilayers.

Computationally, it is now understood that protein-lipid interactions are recruited and protein-specific.^20^ Experimentally, it has been seen that some protein conformations are inaccessible when reconstituted within a given membrane composition, a finding further consolidated by the ability of detergents to facilitate or deter membrane protein crystal structures from certain conformational states.^21^ Performing long timescale simulations studies with realistic membranes can serve as a starting point for improved understanding. Such simulations present an opportunity to expand upon current computational knowledge of protein-lipid interactions and enhance insights to lipid-dependent membrane protein conformational ensembles.

Herein, a realistic plant plasma membrane construct is chosen as a model system to observe the effects of variable membrane complexity on pure membrane properties and membrane protein function. Currently there exists no standard protocol for conducting MD simulations on membrane systems modeling a plant cell; unlike for mammalian systems, whose bilayers are generally assumed to be POPC-rich,^22^ plant membrane systems lack extensive study or even default compositions when simulated via MD. Some plant membrane compositions exist, though have mostly been used for a specific application, such as the thylakoid membrane for photosynthesis studies, and are not asymmetric.^23–27^ While plant species present biological differences, an averaged, generalizable model would facilitate membrane MD study in plants. Furthermore, by establishing a membrane composition with valid asymmetry whose physical properties are robust against size-dependent changes, the effects of membrane realism on molecular simulation timescales can be evaluated. For comparing the effects of membrane composition on protein function, two formulas for plant plasma membrane construction were then implemented: (1) a composition which was purely based off experimental ratios scaling with respect to the membrane size (called “Maximum Complexity”), and (2) a size-independent composition where only the top ten most abundant lipids species – those which had a literature calculated relative abundance greater than 1% - were retained (termed “Top10”). Depending on the system size, our all-atom plant plasma membranes are represented by the asymmetrical arrangement of 9, 10, 15, or 16 different lipid species. Such chemical diversity is rarely seen in all-atom MD approaches to membrane modeling. Model bilayers containing 10 or more different lipid species exhibited more or less equivalent physical properties.

To this end, the *apo* form of OsSWEET2b, a rice sugar transporter, was inserted into each membrane type. The complete transport cycle of *apo* and *holo* variants of OsSWEET2b had previously been elucidated in a pure POPC membrane,^28^ allowing for a direct comparison to evaluate the differences brought by introduction of a native membrane composition. Between the two membrane constructs, OsSWEET2b *apo* transport cycles were capitulated across an aggregate of 61 and 91 *μ*s using Maximum Complexity and Top10 membranes, respectively. Markov state models (MSMs) were used to remove sampling bias and interpret aggregate data from many short simulations into an ensemble view for characterizing longer timescale behaviors (i.e., OsSWEET2b gating dynamics). Thus, the use of MSMs helps in better estimating the free energy barriers between conformationally distinct gating states. While the establishment of a realistic plant plasma membrane is intended for enabling future general simulation applications in plant-based systems, our results concerning transporter dynamics are intended to serve as a reference for MD practitioners considering how complex to make their own membrane models when studying membrane protein function. Ultimately, we advocate for constructing the most realistic membranes when lipidomic data is not limiting. Even in cases where simplification of membrane complexity is unavoidable, we present readers with discussion of some practical considerations when modeling membrane protein function.

## METHODS

### Curation of literature for plant plasma membrane construction

An extensive literature search was performed to determine the lipid composition of a realistic plant plasma membrane representing a generalized plant cell. Curated data including relative percentages of sterol and lipid headgroup composition were averaged regardless of plant species and tissues sampled. Fatty acid chain distributions for each headgroup were estimated only from sources where mass spectrometry information was provided on both the *sn-1* and *sn-2* acyl chains.^29–43^ To make this membrane as realistic as possible, we wanted to account for leaflet asymmetry. In terms of asymmetric lipid distribution across different leaflets within the plant plasma membrane, literature is limited; specifically, it has been reported that the distributions of lipid species across the plasma membrane upper:lower leaflets are 35:65 and 70:30 for phospholipids and sterols, respectively.^44^ However, to the best of our knowledge, no such data is provided on the specific distributions of phospholipid headgroups and acyl chain types across the plant plasma membrane leaflets. We proceeded under the assumption that the headgroup and acyl chain asymmetries seen across the outer and inner membranes of chloroplasts and mitochondria would be similar to that for the plant plasma membrane. From our literature search,^45–58^ we compiled the data of lipid species distributions across the outer and inner membranes of chloroplasts and plant mitochondria to determine relative ratios for lipid species with different acyl chain lengths and degrees of unsaturation within each headgroup type.

### Membrane and membrane protein system assembly

Pure membrane constructs were made with the designated compositions in the CHARMM-GUI Membrane Builder webtool.^10^ Pure membrane constructs were sized at 64, 128, 256, and 512 total lipids. These system sizes were chosen to reflect the necessary membrane sizes needed for common MD applications (e.g., pure membrane property calculations, small-molecule membrane interaction studies, membrane protein structurefunction capture). The largest system size which could be made using CHARMM-GUI was 512 lipids; the larger sterol content resulted in lipid tail-ring penetration issues which could not be overcome when using CHARMM-GUI at larger sizes. For OsSWEET2b simulations, select structures representing previously published inward-facing (IF), outward-facing (OF), occluded (OC), and hourglass (HG) conformations were used for system construction.^28^ One of each of these four protein structures was then submitted to the Membrane Builder webtool for membrane building using the “Maximum Complexity” and “Top10” compositions at a membrane size of 256 total lipids. Addition of all water molecules for each system was then repacked using PACKMOL 18.169.^59^

### System parameterization

CHARMM36 protein and lipid forcefields were applied to each system in the psfgen VMD-plugin,^60^ where protein protonation states were determined using the PDB2PQR (PROPKA) server.^61^ CHARMM-formatted topology and structure files for pure membrane systems over 100,000 atoms in size were generated using the TopoTools plugin to output final structure files in a format other than the pdb format, as pdb files cannot be written to number more than 99,999 atoms.^62^ The default TIPS3P water model for CHARMM was used for all simulations. Potassium and chloride ions were added using the autoionize package to neutralize the systems. All systems prepared through VMD were then converted from CHARMM to AMBER format using the CHAMBER within the ParmEd toolkit to enable parallel computing capabilities in AMBER.^63^ During conversion, all membrane protein systems were subject to hydrogen mass repartitioning (HMR) in order to allow faster timesteps for simulation, which would help with capturing slower kinetic processes within single trajectories.^64^ Pure membrane simulations were not performed using HMR for simplifying characterization work.^65^

### Simulation details – Pure membrane simulations

All simulations were performed by using classical molecular dynamics (MD) in the AMBER18 software package.^66^ For pure membrane simulations, minimization was performed for 50000 cycles, where steepest descent was used for the first 5000 and then conjugate gradient for the remaining 45000 cycles. No positional restraints were employed. Each system was then heated for 2 ns from 0 to 10 K in an NVT ensemble. Each system was then heated at 10K in an NPT ensemble for 2 ns. NPT heating was then increased to 300K for 2 ns. All prior heating stages were run using the `sander` CPU simulation code. In order to ease box dimension changes prior to using the `pmemd.cuda` GPU simulation code, a 5 ns NPT hold was performed at 300K using the `pmemd` CPU simulation code. Following the 5ns hold, all systems then underwent equilibration and production runs. Pure membrane simulations used semiisotropic pressure scaling and were run with a 2fs timestep. A Berendsen thermostat and barostat were used throughout heating steps, where the pressure was maintained at 1 bar.^67^ A Langevin thermostat and Monte Carlo barostat were then implemented for temperature and pressure maintenance in all equilibration runs, respectively.^68,69^ A Langevin collision frequency of 2 ps^−1^ was used, while default settings for the Monte Carlo barostat were kept. Equilibration and production runs proceeded using the `pmemd.cuda` GPU simulation code except for the 64-sized lipid membranes for reasons discussed in the main text, which were run using `sander`. The SHAKE algorithm was applied to all stages of simulation initialization except for minimization,^70^ while the Particle Mesh Ewald method was used for treating long-range electrostatics at a 12 Å cutoff in correspondence with the CHARMM36 lipid forcefields.^71,72^ Pure membrane simulations were equilibrated for 1 *μ*s and in triplicate, where the frame save rate was set to every 50000 frames. Both Maximum Complexity and Top10 models appeared to equilibrate within 400 ns.

### Simulation details – Membrane protein simulations

Membrane protein simulations were minimized in a similar fashion to the pure membrane simulations, although positional restraints at strengths of 5 kcal/mol were applied to all backbone atoms. Because of HMR, all membrane protein simulations were run with a 4fs timestep. Identical heating protocols were used for membrane protein simulations except for the following changes. Anisotropic pressure scaling was implemented throughout and a Berendsen thermostat and barostat were used for equilibration and production runs. Despite their widespread accessibility in MD software, it has not been recommended to use the Berendsen thermostat or barostat for production runs unless comparing results to already published work which had used them.^73^ Accordingly, we used a Berendsen thermostat and barostat to compare the effects of increased membrane complexity on the gating dynamics of OsSWEET2b.^28^ All atom restraints were removed from membrane protein simulations during the 5 ns holding period. OsSWEET2b simulations were equilibrated for 50 ns. Based on expected timescales, OsSWEET2b production runs ran for 25 *μ*s, where only coordinates and not velocities were read from restart files. Velocities are purposefully reinitialized in adaptive sampling in order to achieve greater randomized seeding, and in turn, more efficient sampling over the desired conformational space. Frame recording rates were set for every 50000 steps.

### Adaptive sampling protocols

Pure membrane simulations were run using continuous MD while membrane protein simulations were run using adaptive sampling-based MD. In adaptive sampling regimes, any state recorded along a trajectory can be used as the starting point to seed a new trajectory.^74–77^ A k-means clustering of trajectory data based on some reaction coordinate allows for the potential states used for seeding new trajectories to be representative of the entire round of data collected.^78^ For production trajectories run from each starting OsSWEET2b conformer within each membrane construct, the top 50 structures from the least populated clusters were used to reseed new adaptive rounds of simulation. For *apo* simulations, distances between gating residues (intracellular gating residues Phe42 and Phe164; extracellular gating residues Arg69 and Asp189) were used as the reaction coordinates for clustering. Systematic adaptive sampling was performed until the gating landscape for OsSWEET2b gating dynamics was filled or completed. All distance measurements between atoms were conducted using MDTraj 1.9.3.^79^ Clustered states and trajectories were visualized using VMD 1.9.3, while aggregate data collected were visualized as projected landscapes using Matplotlib 3.2.0.^80^

### Pure membrane simulation analyses – validation of physicality among constructs

The physical correctness of asymmetric membrane simulations can be assessed in two different ways, where either approach attempts to evaluate to what extent lateral pressure distributions differ between leaflets. If the difference in lateral pressure experienced by either leaflet is too great, then the pressure experienced by one leaflet will dominate the dynamics experienced by the bilayer as a whole. One method to clarifying satisfactory lateral pressure differences between leaflets is to compare the area-per-lipid (APL) mismatch between the two leaflets of the asymmetric construct.^11^ As long as APL mismatch is between 5 and 10%, the impacts of asymmetry on bilayer properties are expected to be mild, where the best-case scenario is to have an APL mismatch between 0 and 5%. Another approach is to use the upper and lower leaflet compositions of the desired asymmetric leaflet membrane to build two symmetric membrane constructs.^22^ So long as the membrane-spanning box dimensions of the symmetric systems - in this case, the XY dimensions – demonstrate a similar box area difference from the asymmetric simulations, the defined leaflet asymmetry can be deemed acceptable. Symmetric membranes were built for each membrane size tested, prepared as described above, and with production runs totaling for 50 ns. The box dimensions from these trial simulations were then compared with the first 50 ns of asymmetric membrane simulations.

### Pure membrane simulation analyses – bulk and headgroup-specific physical property measurements

Pure membrane simulations were analyzed for bulk physical properties using two software: Membrainy and SuAVE.^81,82^ The following physical properties were calculated using these software: area-per-lipid (APL), entropy, gel fluidity, membrane thickness, membrane curvature order parameter, membrane curvature angle, lipid *S*_CD_ order parameters, headgroup axial tilt angle, and headgroup lateral tilt angle. Electron density calculations were performed using the CPPTRAJ module found in AMBER.^83^ Membrainy input files were generated by converting the AMBER-formatted trajectory and topology files to GROMACS formats using GROMACS 2020.^84^ SuAVE input coordinates were generated from trajectories using VMD and MDTraj.

### Membrainy software updates

Membrainy was updated to allow new lipids to be parameterized via a user-created external lipid library (i.e., accept custom lipid forcefields).^81^ The leaflet detection method was also modified for improved efficiency and to allow for a vector-based solution to better account for leaflet undulation when discriminating between leaflets for calculations. In this leaflet-identification approach, the vectors are created by using the phosphorous atom and the last carbon atom of each lipid tail. For lipids with two tails, the average vector orientation was taken. Sterols are treated as having only one tail. Vectors that point up indicate an individual lipid getting placed in the upper leaflet, with vectors pointing down indicating lower leaflet placement. An updated version, Membrainy 2020, capable of performing all calculations done in this manuscript is available online (http://www.membrainy.net/) and is compatible with Gromacs-2020.^84^

### OsSWEET2b Markov state model (MSM) construction and validation

All stages of MSM construction were performed using PyEMMA software.^85^ Feature selection for OsSWEET2b MSM construction followed the same protocol as employed in previous studies on membrane proteins.^28,86–88^ Maintaining the same choice of input features enables a direct comparison to the previous OsSWEET2b work done in a pure POPC bilayer. An iterative process of hyperparameter selection for optimization of the maximum VAMP score was performed.^89^ The number of clusters tested for data clustering varied from 100 to 1000 clusters, increasing by increments of 50. The number of time-Independent Component (tIC) were varied from 1 to 9. The time-independent component analysis (tICA) lag time varied from 5, 6, 7, 10 and 12 ns. The optimal combination of hyperparameters for OsSWEET2b Scale MSM construction was 9 tICs, 1000 clusters, and a tICA lag time of 12 ns. For constructing MSM of OsSWEET2b in the Top10 membrane model, the optimal combination of hyperparameters was 9 tICs, 650 clusters, and a tICA lag time of 7 ns. Provided the nine residue pair distances used for featurization, the slowest timescale process for OsSWEET2b dynamics in either membrane converged at an MSM lag time of 9 ns. Accordingly, final MSMs were constructed using the optimal hyperparameters with an MSM lag time of 9 ns.

### Adaptive sampling error analysis

The data ensemble containing nine residue pair distances used for MSM featurization were replicated into ten slices. Each of these slices were comprised of a random selection of 80% of the overall data. MSMs were constructed off each of these data subsets using the same combination of hyperparameters as the entire dataset. With the gating dynamics represented as a 2D histogram plotting extracellular versus intracellular gating distances, the average error for each bin among each of the data slices was calculated as previously shown.^28^

### OsSWEET2b annular shell analyses

The linear interaction energy between OsSWEET2b residues and lipid species was calculated in CPPTRAJ. Snapshots were rendered using Chimera 1.14.^90^

## RESULTS AND DISCUSSION

### Establishing a realistic plant plasma membrane composition

Based on the literature survey, the plant plasma membrane presents an ~48:52 split between sterols and phospholipids (PL) across the entire bilayer. However, the upper leaflet has a sterol:PL split of 65:35.^44^ Meanwhile, the lower leaflet exhibits a sterol:PL split of 30:70,^44^ which is consistent with the “positive-inside” rule, as more electropositive phospholipid headgroups are present along the intracellular leaflet to counter cytosolic electronegativity (Figure 1).^91–93^ In order of abundance, the plant plasma membrane is comprised of SITO (~32%), STIG (~16%), PLPE (~13%), PLPC (~12%), DLiPC and DLiPE (~5%), LLPC and PLPG (~4%), LLPE (~3%), POPC (~1%), and DSPC, DOPC, POPE, SLPE, POPG and DPPG (< 1%). While PLs represent the aggregate bulk of the plant plasma membrane, the most abundant individual lipid species are sterols, with SITO and STIG at ~32% and ~16%, respectively (Figure 2).

**Figure 1.**
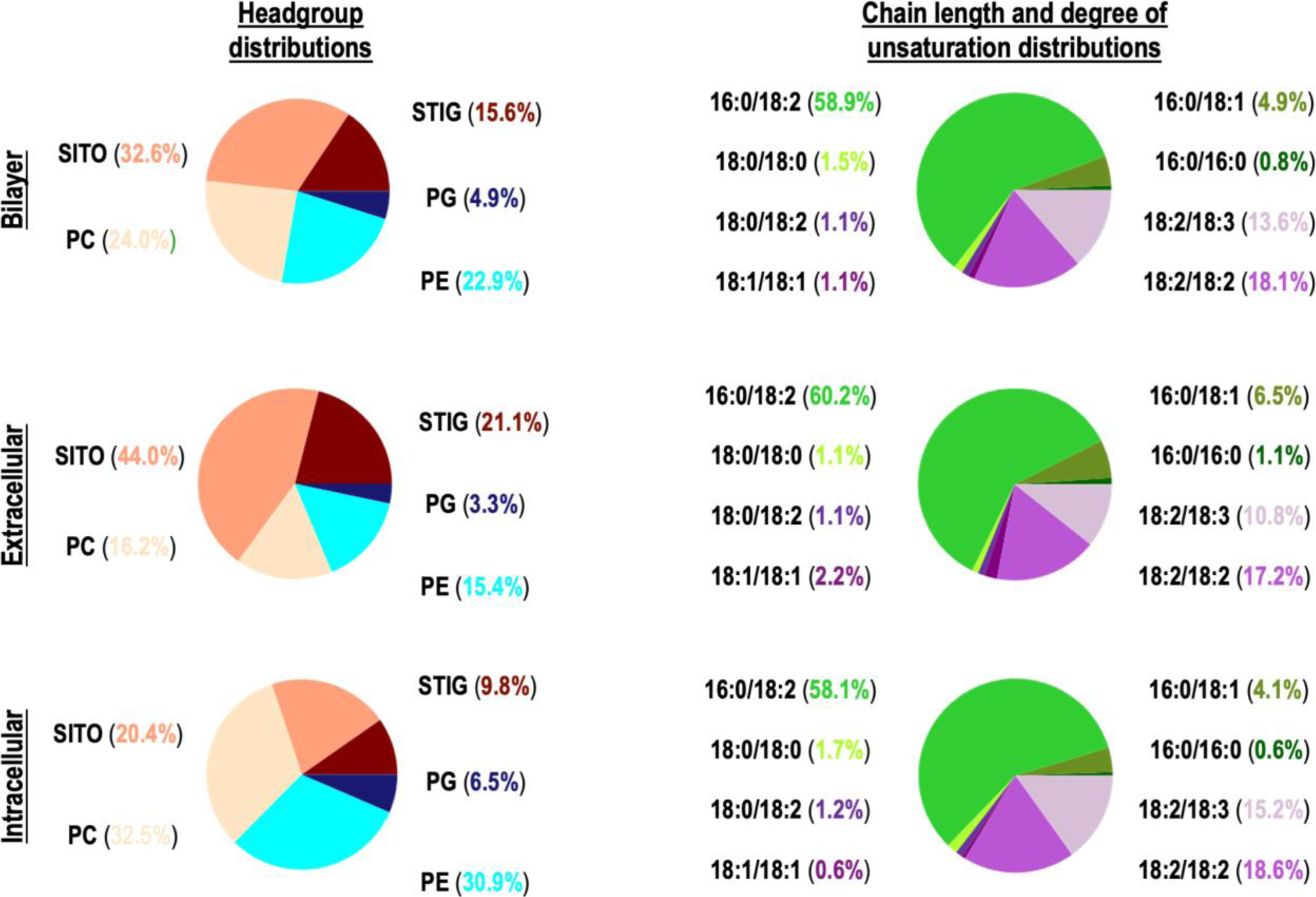
Distribution of different lipid headgroups and degrees of tail unsaturation in model membranes constructed following a Maximum Complexity recipe.

**Figure 2.**
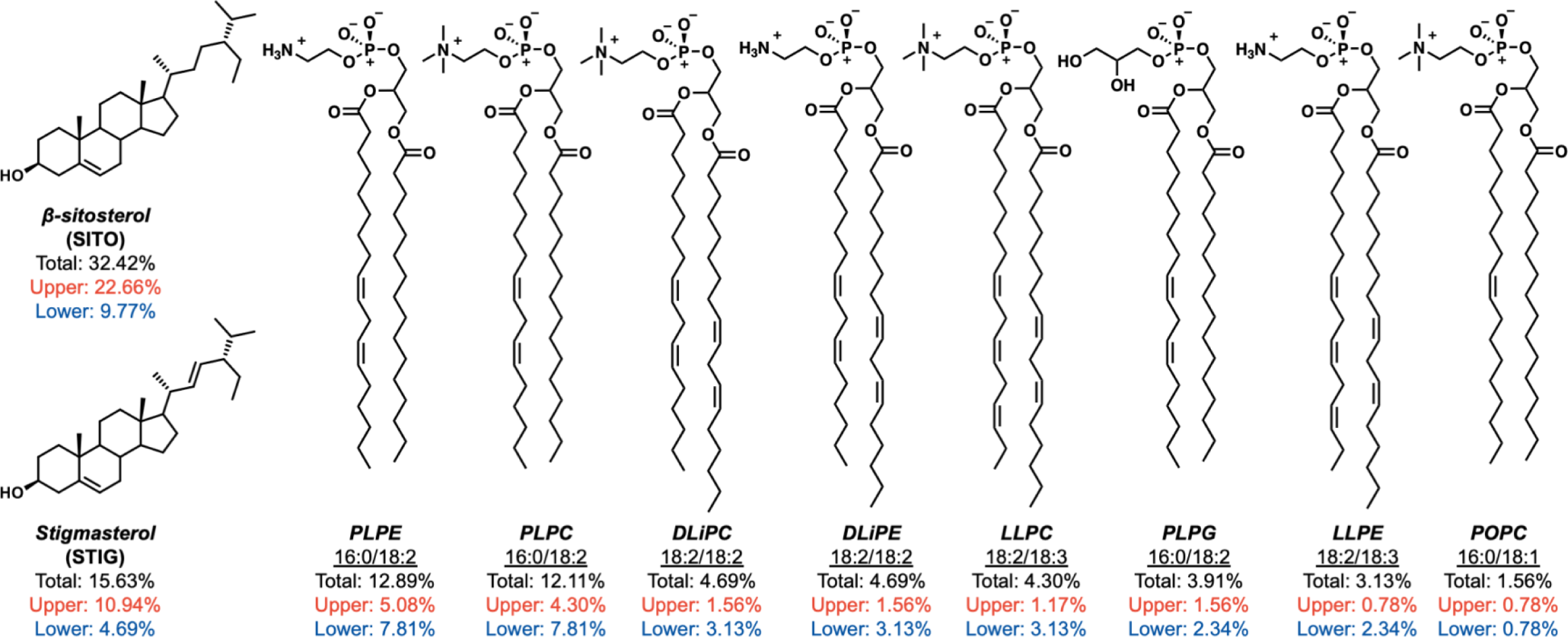
Leaflet-specific percentage of different lipids present in a Maximum Complexity membrane construct. Ratios presented are based off those seen in the 256-lipid construct.

Such sterol abundance has been seen in other membrane compositions.^94^ The most abundant individual PLs are PLPE (~14%) and PLPC (~12%) (Figure 2). Still, PC headgroup lipids account for the majority of PLs, followed by PE and PG. Regardless of leaflet, PLs predominantly exhibit 16:0/18:2 unsaturation (~60%), followed by 18:2/18:2 (~18%). Taken together, ~91% of PLs exhibit 18:2 unsaturation along one of their acyl chains. This is a far departure from simple mammalian membrane models which are often simulated using just POPC (16:0/18:1), serving as a reminder that “default” membrane practices for one organism type cannot be assumed to be appropriate for modeling another organism. For comparison, POPC represents ~1% of the plant plasma membrane composition, although the type of 18:1 unsaturation present in POPC-like lipids accounts for ~5% of all lipid unsaturation (Figure 1). The exact ratios for Maximum Complexity lipid bilayers for each size constructed (64, 128, 256, and 512 total lipids) are supplied in Tables S1-S5.

Reducing the Maximum Complexity membranes to that of a Top10 composition, where any lipid species present under 1% relative abundance is removed from consideration, causes the relative abundances of the remaining lipid species (SITO, STIG, PLPE, PLPC, DLiPC, DLiPE, LLPC, PLPG, LLPE, and POPC) to fluctuate slightly. Removal of just five lipid species with < 1% relative abundance causes the remaining ten species to be represented in greater abundance, as the MD system membrane size is the same but now with fewer types of lipids. Accordingly, this exacerbates leaflet-specific asymmetries as lipid counts become redistributed (i.e., the presence of slightly less sterols in the upper leaflet, or slightly more PLs in the lower leaflet, than would typically occur in nature). For the sterols, the disparity in relative abundance between the two leaflets decreases, where more sterols are redistributed to the lower leaflet. The reallocation of trace lipid counts to other lipids within their same headgroups results in relative increases for all PLs except for LLPE and POPC. PLPE content is increased by ~3%, making it now the second most abundant lipid species above STIG (Figure S1).

### Pressure-related analyses indicate membrane physicality

To evaluate the generalizability of this informatically-driven Maximum Complexity plant plasma membrane composition and its simplified Top10 version, common membrane physical properties (box area, area per lipid, entropy, gel fluidity, thickness, curvature, acyl chain order parameters, headgroup orientation, electron density) were calculated.

Asymmetric membranes inherently experience differing lateral pressure profiles due to the different number of lipids present in each leaflet. Numerous developments by the field have been made to mitigate the damage to membrane physicality as a result of inherent differences between the leaflets of an asymmetric membrane. The generation of optimized and adjusted APL profiles for a given lipid species and zeroed lateral tension settings are some ways to help.^8–12^ Ultimately, asymmetric membrane physicality (i.e., how distinct lateral pressure profiles are for each leaflet) needs to be verified. Short simulations using symmetric bilayers for each leaflet of each membrane construct can be used to retrieve an MD box size specific to that leaflet pairing. For example, the upper leaflet from a Maximum Complexity construct was used to create a symmetric bilayer. This is critical for validating asymmetric bilayers, as the individual leaflets must have stable individual pressure constraints. The box size for a membrane system is a valuable metric for evaluating membrane construct stability. Given that the membrane bilayer spans the entire MD box in the X and Y dimensions, box size fluctuations give an estimate as to how the bilayer responds to the barostat. So long as the calculated box sizes between the symmetric leaflets are not too different from each other and the final asymmetric construct, then it can be viewed as reasonable for the specified leaflet pairing to comprise a single bilayer.^22^ Another validation approach is to compare the percentage difference in APL between the two leaflets of an asymmetric bilayer.^11^ Similar to box size, APL serves as another measure to evaluate how the bilayer is responding to pressure. When APL converges, a membrane is often said to have equilibrated. According to this approach developed by the Klauda and Im groups in 2015, the ideal APL mismatch between leaflets of an asymmetric membrane should be between 0 and 5%.^11^ Yet membranes demonstrating 5 to 10% mismatch only demonstrate mild effects from asymmetry, the extents of which are less noticeable in membranes with more unsaturated tails.

A general rule of thumb is to assume that membranes containing more than one type of lipid species requires 100 ns of equilibration for each of the different lipid species present.^95^ From APL analyses it appears that each of the membrane constructs equilibrated after 400 ns of simulation, although the APL for each 64-sized bilayer leaflet appear to be distinct from the other membranes’ (Figure 3). APL mismatch analysis shows the 128- and 256-sized Maximum Complexity membrane constructs with 6.06% mismatch between leaflets (Table 1). Meanwhile, the Maximum Complexity constructs containing 64 and 512 lipids have 11.76% and 7.52% leaflet APL mismatch, respectively (Table 1). When comparing the symmetric constructs for each asymmetric leaflet to the asymmetric bilayer, the leaflet area differences are all below 5%, except for the 64-lipid lower leaflet construct (Table 2).

**Figure 3.**
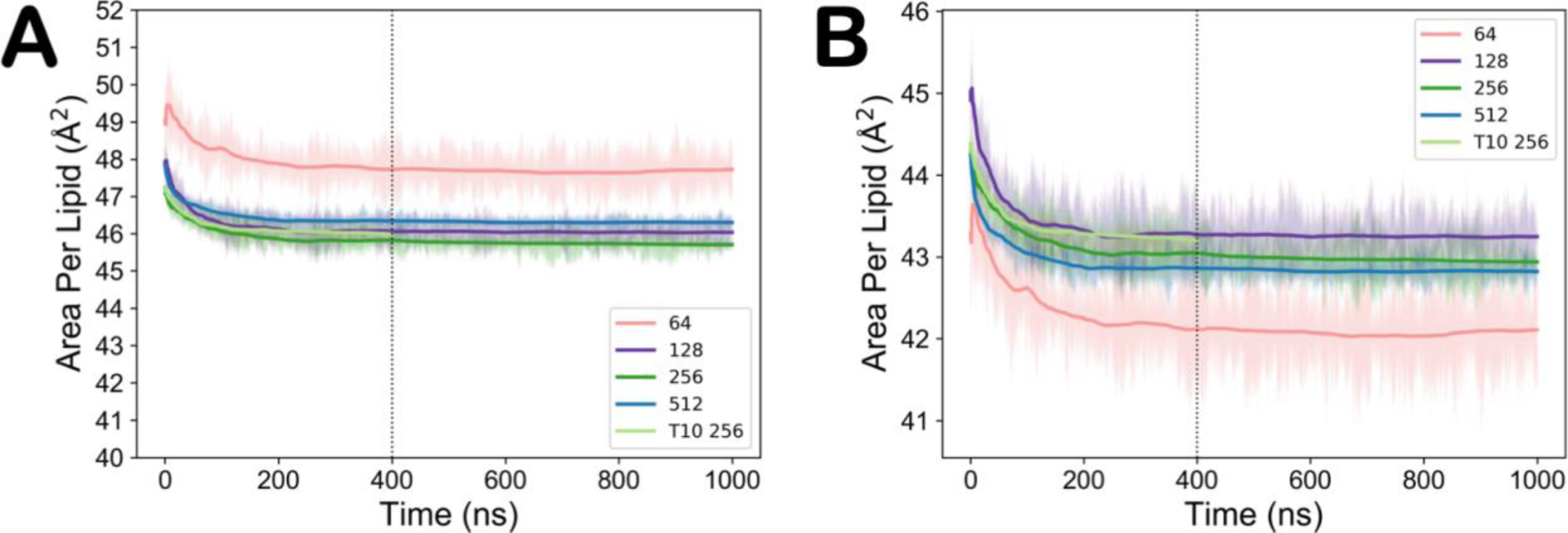
Area per lipid (APL) values for the (A) lower and (B) upper leaflet of different sized membrane constructs.

**Table 1.** APL and APL % mismatch between leaflets of membrane constructs

| Systems | AVG APL<br>Lower Leaflet | AVG APL<br>Upper Leaflet | AVG APL<br>% mismatch | SEM APL<br>% mismatch |
| --- | --- | --- | --- | --- |
| Maximum Complexity 64 | 47.72 ± 0.26 | 42.11 ± 0.23 | 11.76 | 6.84E-14 |
| Maximum Complexity 128 | 46.04 ± 0.14 | 43.25 ± 0.13 | 6.06 | 4.23E-14 |
| Maximum Complexity 256 | 45.71 ± 0.17 | 42.92 ± 0.16 | 6.06 | 1.29E-13 |
| Top10 256 | 45.98 ± 0.03 | 42.82 ± 0.03 | 6.06 | 4.62E-14 |
| Maximum Complexity 512 | 46.30 ± 0.04 | 42.82 ± 0.04 | 7.52 | 2.92E-13 |

**Table 2.** Leaflet-specific pressure tests. Comparing box area difference between symmetric leaflet constructs and final asymmetric bilayers.

| Systems | Symmetric System Data |  |
| --- | --- | --- |
|  | Lower Leaflet Area Difference (%) | Upper Leaflet Area Difference (%) |
| Maximum Complexity 64 | 5.86 | 2.59 |
| Maximum Complexity 128 | 5.03 | 4.7 |
| Maximum Complexity 256 | 4.52 | 4.14 |
| Top10 256 | 3.28 | 0.71 |
| Maximum Complexity 512 | 4.53 | 3.17 |

The leaflet area deviations from the asymmetric bilayer are most similar between symmetric Maximum Complexity constructs containing 128 and 256 lipids, which is in agreement with the APL analyses. Thus, when generating plant plasma membranes for simulation, asymmetric bilayers between 128 and 256 lipids in size would be expected to be least affected by lateral pressure differences caused by asymmetry, followed by the 512-sized membrane. The APL mismatch for the 64-sized bilayer suggests that lateral pressure differences experienced by either leaflet would compromise simulation integrity, likely leading to unrealistic or unphysical representation of the membrane. No 512-sized Top10 construct which arrived at the equilibration stage of simulation made from the given lipid could pass either of these pressure tests because of leaflet-specific asymmetry being too different. Thus, the asymmetry recipe applied to this simplified Top10 construct was only valid for a 256-sized bilayer.

### Physical property analyses demonstrate robust compositional recipes

The remaining bulk property analyses included here are APL, entropy, gel fluidity, thickness, and curvature. Mixing entropy indicates the probability that a lipid of one type will have a chemically different lipid as its neighbor. The higher the lipid type mixing entropy, the more mixed the membrane.^96^ For each of the leaflets, entropy analyses follow a size-dependent trend. Given that compositional complexity scales with membrane size when following the Maximum Complexity recipe, it is expected that larger membranes would have higher entropy or mixing probabilities (Figure 4). However, the 256-sized Top10 construct violates this relationship, where its upper leaflet has a higher entropy than the 512-sized membrane. Overall upper leaflet entropy values appear to be more similar between each of the membrane constructs, suggesting that their shared sterol-rich profiles may underlie similarities in lipid organization. Like upper leaflet entropy, 256-sized Top10 construct bilayer entropy matches that for the 512-sized Maximum Complexity construct.

**Figure 4.**
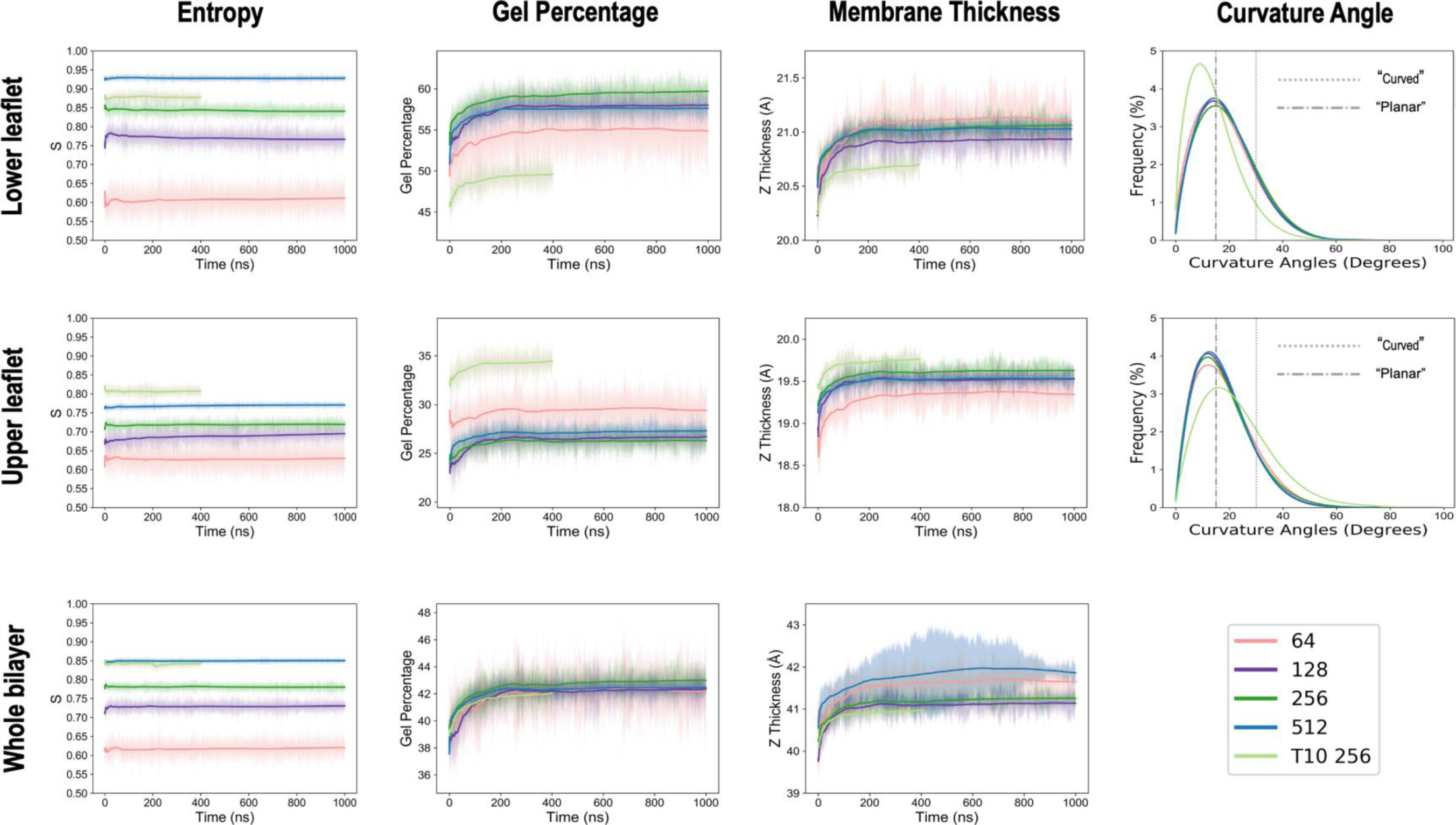
Membrane construct physical property analyses demonstrate robustness of compositional recipe.

Gel fluidity measures the linearity of the lipid tails, which can provide insights into the bulk ordering of the membrane.^81^ Here, all membrane constructs demonstrate the same bilayer properties, but the fluidity of each of the leaflets differs. For either leaflet, the 64-sized Maximum Complexity and 256-sized Top10 constructs appear to have distinct distributions of gel fluidity percentage from the remaining construct sizes, where the Top10 construct’s lower leaflet is less and its upper leaflet is more fluid than the Maximum Complexity constructs. Membrane thickness calculations follow similar, albeit less pronounced per leaflet, trends as gel fluidity (Figure 4). Membrane curvature calculations lastly show that regardless of size, all Maximum Complexity membranes demonstrate the same curvature distribution, peaking at a curvature angle of 18° (indicative of a planar surface).^82^ Meanwhile, the Top10 membrane construct differs, where the lower leaflet is slightly more planar and the upper leaflet more curved (Figure 4).

Headgroup-specific properties measured include acyl chain order parameters, headgroup orientation, and electron density. Surprisingly, when comparing properties between lipid species which are shared between each of the membrane constructs, all of these results are nearly equivalent or within error (Figures S2-S13). It appears that headgroup-specific properties are not as influenced by the relative system complexity nor size. On the other hand, bulk membrane properties differ only slightly between a few membrane constructs. The 256-sized Top10 construct appears to have altered distributions from the 256-sized Maximum Complexity membrane likely because of composition-dependent effects. Again, the major differences seen in the 256-sized Top10 construct were the relative redistribution of sterols to the lower leaflet and the amplification of 18:2 lipid species across the bilayer. It is possible that slight alteration of relative lipid abundances could impact the initial packing schemes within CHARMM-GUI. As for the 64-sized Maximum Complexity construct, differences in bulk properties can be accredited to a number of factors. Pressure tests (Tables 1 and 2) indicate that the asymmetry seen in such a small membrane may not be physically valid. This discrepancy may be exacerbated by an analytical effect from the MD engine. The box-size for 64-sized Maximum Complexity constructs approached the minimum nonbonded cutoff specifications required to perform simulations using Amber software. Thus, 64-sized membranes could only be simulated using the more sensitive CPU-based code, `sander`, which may amplify any size-dependent effects when compared to the systems run using the GPUbased code `pmemd.cuda`.^66^ Aside from the abovementioned constructs, Maximum Complexity constructs sized 128 to 512 lipids appear robust enough to be used for any plant membrane MD application and demonstrate equivalent properties.

### Effect of simplified complexity on membrane transporter dynamics

OsSWEET2b, a vacuolar sugar transporter, was selected for modeling the effects of reducing membrane complexity on membrane protein function. Although OsSWEET2b is not found in the plant plasma membrane, it is suitable to study for two demonstrative purposes. Firstly, the *apo* transport cycle for this protein has previously been elucidated in a symmetric and homogenous POPC membrane.^28^ Simulation of OsSWEET2b enables direct comparison of the influence of using a complex membrane construct versus an overly simplified one. Secondly, detailed lipidomic data providing both acyl chain identities for constituent lipids of a tonoplast (vacuolar) membrane does not exist. Indeed, this lack of high-resolution lipidomic data concerning relative lipid species abundances is a fundamental driver causing MD practitioners to resort to very simple membranes. While exact structure elucidation is missing, plant tonoplast membrane composition has been measured at low resolution with the measurements of headgroup and single acyl chain abundance.^43,50,53,97–107^ Overall, the plant tonoplast offers similar trends in relative headgroup and individual acyl chain representation to suggest that the same lipids would be represented as shown in these plasma membrane constructs. Thus, based on rationale and experimental evidence suggesting enough similarity between tonoplast and plasma membrane compositions, we proceed with simulating the transport cycles of OsSWEET2b.

To evaluate the effect of protein-lipid interactions, it is advisable to generate multiple versions of the same membrane composition with different initial packings.^108^ By using multiple membrane starting structures to generate different membranes with the same overall composition, a greater diversity of potential protein-lipid interactions can be observed, thereby providing greater perspective into the effects of a complex membrane composition.^108^ Such a project design also makes the capture of OsSWEET2b gating dynamics more efficient as it enables greater initial starting structures for seeding adaptive sampling-based trajectories.^74,109^

OsSWEET2b gating dynamics cover roughly the same conformational spread of intracellular and extracellular gating distances when simulated with the either the Maximum Complexity or Top10 membrane constructs. Both landscapes present reasonable sampling of extracellular gating distances ranging between ~4-20 Å and intracellular gating distance ranges of ~5-25 Å (Figure 5). Small deviations between the energetic accessibility of intermediate states of each landscape exist, although the majority of them fall within the 0.2 kcal/mol sampling error (Figure S14). For example, the extent of stabilization for IF states differs between the constructs, where the Top10 construct demonstrates an energetic cost of ~0.6-0.8 kcal/mol versus ~1.2-1.4 kcal/mol seen in the Maximum Complexity construct (state 2 of each landscape), which falls within error (Figure 5). Energetic costs underlying some transitions exceed sampling error. The difference in transition energy from Maximum Complexity states 2→4 and the corresponding Top10 path between states 2→3 exceeds the error limit, exhibiting free energy costs of 2.8 ± 0.2 and 3.5 ± 0.3 kcal/mol, respectively (Figure 5). A similar analysis on the most probable path between Maximum Complexity states 3→5 and the corresponding Top10 states show respective free energy barriers of 1.8 ± 0.2 versus 2.4 ± 0.3 kcal/mol.

**Figure 5.**
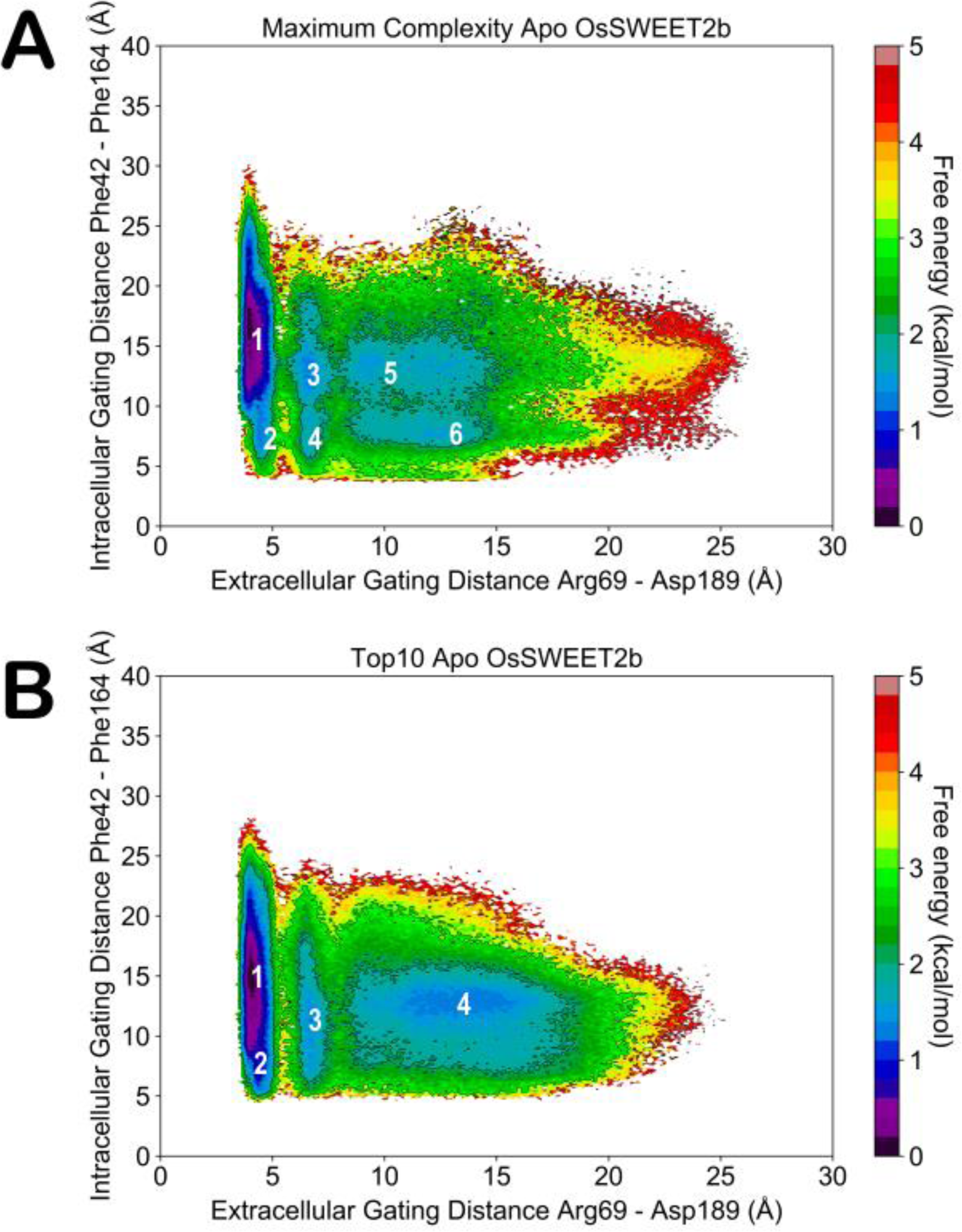
MSM-weighted OsSWEET2b gating dynamics. (A) apo OsSWEET2b simulations done in the 256 Maximum Complexity membrane construct. (B) apo OsSWEET2b simulations done in the 256 Top10 Complexity membrane construct. Key conformational states are numerically labeled.

Aside from more stringent quantitative analyses, visual inspection of the gating landscape shows how the Maximum Complexity membrane construct is able to better resolve microstates seen with respect to IF, extended IF-OC, and extended OF-OC gating states (Figure 5). OsSWEET2b Top10 simulations indicate more coarse and less comparable lumping of OF states, where sampling of a more OF and less OC state is less likely when compared to the Maximum Complexity gating landscape. This coarser representation of Top10 microstates also reflects the slightly higher barriers between intermediate OC states and fully IF or OF states. When compared to the Maximum Complexity construct, the simplification of lipid composition provided by the Top10 construct does provide a smoothening effect to the one-dimensional projections of either the intracellular and extracellular gating distances, particularly when it comes to resolving IF-OC and OF-OC states (Figure S15). The thermodynamic differences between Maximum Complexity and Top10 conformational landscapes are small (~ 1 kcal/mol) but these differences based on lipid composition could lead to significant changes in kinetics and impact conformational sampling.

### Exploring specific protein-lipid interactions with varying membrane complexity

Regardless of which realistic membrane composition was used, *apo* simulations performed using a native versus a purely POPC composition demonstrate overall lower energetic costs for transport (Figure S16). In fact, the *apo* gating dynamics landscape for simulations done in a realistic membrane composition more closely resemble the previously published *holo* gating dynamics performed in a purely POPC membrane composition.^28^ Between the ~1 kcal/mol lower barrier for all transitions and the protein spending less time in higher energy intermediate states, the extent of stabilization provided by a native membrane environment mimics the energetic stability provided by substrate introduction, suggesting that specific interactions between OsSWEET2b and annular shell lipids could be a contributing factor.

An annular shell analysis, which differentiates the lipids in the immediate vicinity of the protein compared to the bulk, was performed on OsSWEET2b states from a higher energy transitionary region containing extracellular gating distances between 5.25-5.75 Å and intracellular gating distances between 13.25-13.75 Å from simulations involving either membrane construct. This region covers a transition between an IF and extended IF-OC conformational state and is one where minor discrepancy in free energy differences between each of the membranes exists. Each of the states were visualized to see which lipids presented themselves within 7 Å of the OsSWEET2b transmembrane-spanning surface. The annular shell compositions provided by the four different system setups for each membrane construct mainly differ in leaflet-dependent manners. Upper-leaflet sterol and PE content, as well as 16:0, 18:1, and 18:2 unsaturation lipid tails differ between annular shells formed in Maximum Complexity and Top10 constructs (Figure S17). Meanwhile the lower leaflet annular shell composition is more or less equivalent minus the presence of 18:0 and 18:1 unsaturated lipid tails (Figure S17). Analyzing snapshots revealed the presence of a putative lipid binding site with interaction energies largely dominated by Tyr168. Defining a “full binder” as a lipid species whose scaffold fits entirely into the cleft formed by residues Ile17, Phe18, Leu24, Val27, Thr28, Tyr168, Leu171, and Leu175, Maximum Complexity membrane setups never pose a sterol as the “full binder” while Top10 membrane setups can (Table S6, Figure S18). Top10 membrane setups can additionally pose sterols as partial binders which are able to interact with Tyr168; this was not seen in any of the Maximum Complexity setups, each of which had more stable interaction energies with a PL by at most ~1.8 kcal/mol (Table S6). It is possible that even slight modification in relative lipid abundance results in altered packing of specific lipid species. For example, redistribution of sterols to the lower leaflet in Top10 system setups could alter the ordering of PLs with large polar headgroups, like PC.^110–113^ Small modifications in bulk lipid ordering could present changes to which conformations of what lipid species would be compatible for annular shell constitution. It could be that altered contacts, such as those seen in this putative lipid binding site, found in different membrane constructs are a result of tiny changes in membrane complexity and lipid species representation. Given some combination of specific membrane composition and protein, such small changes could manifest into energetic differences between the accessibility of some conformational transition paths over others. Again, the specificity of protein-lipid interactions and dependence of initial or achieved annular packings given a bilayer composition offer an element of kinetic control in making conformational sampling feasible.

## CONCLUSIONS

This work establishes the composition of a realistic model plant plasma membrane and utilizes this construct as a model system for examining the effects of realistic membrane composition on long timescale membrane protein dynamics. Hurdles to simulating realistic membranes include lack of lipidomic data and trial-and-error workflows to appropriately account for leaflet asymmetry. After establishing the robust properties of the Maximum Complexity membrane composition, we show that even the removal of trace lipid species can alter the free energy barriers of intermediate conformational states for a given membrane protein. Simulations performed herein using native lipid bilayers provide easier access to intermediate states and may give faster kinetics than a nonnative lipid bilayer. The use of an appropriate membrane appears to become crucial when sampling for rare events and optimizing computational efficiency for membrane protein simulations. More importantly, a more native lipid bilayer produces different results than overly simplified homogeneous lipid bilayers, providing greater confidence and accuracy in the resolution of the membrane protein conformations sampled.

Either for simplicity’s sake or out of necessity, MD practitioners are too often without proper lipidomic data (i.e., mass spectrometry results showing relative abundances for both lipid acyl chains in a leaflet-specific manner) when assembling membrane lipid bilayers for their own studies, ultimately resorting to “default” membrane compositions. Simplistic lipid bilayers for one cell type may be unacceptable for another organism; only 1% of plant plasma membranes are comprised of POPC, which may not even be represented in membrane systems of smaller scale. True, membrane proteins can still function so long as the surrounding lipid environment provides headgroups and acyl chains which are reminiscent of the native membrane composition,^114,115^ but function in a nonnative setting may not be optimal. From the perspective of simulation, protein insertion into an incorrect membrane could result in misrepresentation of state probability, or the generation of a different landscape altogether. In this sense, the ability of an incorrect membrane environment to eschew an energy landscape is similar to that imposed by mutation.

With the increasing availability of computational power and the demand for larger, more accurately represented lipid bilayers, traditional simplistic approaches will no doubt be pushed aside for more complex systems, allowing true representations of realistic membrane-protein dynamics. OsSWEET2b simulations performed in a realistic plant membrane mirrored the dynamics seen in *holo* simulations performed using purely POPC membranes, emphasizing how lipid species can behave as ligands themselves (Figure S16). General protein-lipid binding motifs have been shown,^116,117^ but the lack of consistent realistic membrane modeling when it comes to membrane protein simulations currently leaves this knowledge unexploited. Introducing a large diversity of lipid species into a finite-sized MD membrane system also complicates the annular shell composition for a given protein. While additional membrane simulation approaches have been suggested for modeling the effect of protein-lipid interactions,^108^ we recommend that attention be paid to packing procedures chosen for complex membrane assembly.^118^ Replicating complex membrane assembly is one way to better account for how lipid diversity interfaces with membrane protein function via varied annular shell composition. Small differences in annular shell composition between the two membrane constructs accompanied OsSWEET2b gating dynamics which could not equally access the same intermediate states. It is possible that small changes in very complex membrane compositions could result in altered lipid organization, which may directly impact modeled membrane protein function and dynamics. For instance, hydrophobic mismatch – when the thicknesses of the transmembrane spanning region of a membrane protein and the surrounding lipid bilayer are incompatible – can result in energetic penalties which either restrict or promote certain protein conformational changes.^114,119,120^ Thus even small manipulations in lipid bilayer composition could directly impact membrane protein function by affecting the probability for the optimal annular shell composition to be assembled for a given protein conformation during bilayer construction. This may explain why explain why some intermediate states are not as accessible for OsSWEET2b while embedded in the Top10 versus the Maximum Complexity construct. Furthermore, optimal protein-lipid interactions can be completely lost when using traditional simplistic lipid bilayers (e.g., just POPC) where hydrophobic mismatch penalties could be maximized.

A caveat to these findings is that comparative membrane modeling was only performed on one protein, OsSWEET2b. Here we show that removal of minor lipid species with relative abundances < 1% do indeed alter the accessibility of some intermediate conformational states when using a realistic membrane. Still, more studies on other membrane proteins are required to obtain a generalizable understanding of how non-optimal versus realistic membrane compositions affect the thermodynamics and kinetics of protein dynamics. Current understanding of how to universally approach and contextualize protein-lipid interactions in MD simulations is as unrefined as the present implementation of realistic membranes in biomolecular simulations. The study of complex membrane composition and membrane protein function often appear as mutually exclusive works. Increased prevalence of realistic membranes in membrane protein function studies will ease these two simulation communities to become more conjunctive. As the modeling of more realistic membranes for protein function simulations becomes more standard, deeper simulation analyses will become more routine and facilitate improved translation of *in silico* findings to *in vitro* or *vivo*. With the intent of being used as a modeling practices resource, the results of this work will aid in the encouragement of realistic membrane simulations and increased research output in plant-based membrane MD applications.

## ASSOCIATED CONTENT

### Supporting Information

Simulation results: membrane compositions for specified construct sizes, order parameters, headgroup orientations, electron density, MSM sampling error, 1-dimensional gating landscapes, comparative landscape with previously published work on OsSWEET2b embedded in a POPC membrane, linear interaction energies with residues in putative lipid binding site, annular shell snapshots.

Additional files including raw data, analysis and plotting code, as well as downloadable membrane structures and trajectories, are available free of charge at: https://uofi.box.com/s/6s4jhq5j5pifxofr6ptdvtaply40sslo

## AUTHOR INFORMATION

### Author Contributions

ATW and DS conceived the project. DS supervised the project. ATW gathered literature for membrane design and performed simulations. MC customized Membrainy code for complex membrane analyses. ATW and DS analyzed the data. ATW wrote the manuscript with input from DS and MC. All authors have given approval to the final version of the manuscript.

## ACKNOWLEDGMENT

This research is part of the Blue Waters sustained-petascale computing project, which is supported by the National Science Foundation (awards OCI-0725070 and ACI-1238993) the State of Illinois, and as of December, 2019, the National Geospatial-Intelligence Agency. Blue Waters is a joint effort of the University of Illinois and its National Center for Supercomputing Allocations. We thank Balaji Selvam for providing MSM-weighted OsSWEET2b starting structures and landscapes from a previous work for more amenable comparisons. We also thank Matthew Chan and Soumajit Dutta for useful input during the preparation of this manuscript.

## ABBREVIATIONS

MSM: Markov state models
MD: molecular dynamics
PDB: protein data bank
GPU: graphical processing unit
CPU: central processing unit
HMR: hydrogen mass repartitioning
APL: area-per-lipid
tIC: time-independent component
tICA: time-independent component analysis
PL: phospholipid
SITO: β-sitosterol
STIG: stigmasterol
PLPE: 1-palmitoyl-2-linoleoyl-*sn*-glycero-3-phosphoethanolamine
PLPC: 1-palmitoyl-2-linoleoyl-*sn*-glycero-3-phosphocholine
DLiPC: 1,2-dilinoleoyl-*sn*-glycero-3-phosphocholine
DLiPE: 1,2-dilinoleoyl-*sn*-glycero-3-phosphoethanolamine
LLPC: 1-linoleoyl-2-linolenoyl-*sn*-glycero-3-phosphocholine
PLPG: 1-palmitoyl-2-linoleoyl-*sn*-glycero-3-phosphatidylglycerol
LLPE: 1-linoleoyl-2-linolenoyl-*sn*-glycero-3-phosphoethanolamine
POPC: 1-palmitoyl-2-oleoyl-*sn*-glycero-3-phosphocholine
DSPC: 1,2-distearoyl-*sn*-glycero-3-phosphocholine
DOPC: 1,2-dioleoyl-*sn*-glycero-3-phosphocholine
POPE: 1-palmitoyl-2-oleoyl-*sn*-glycero-3-phosphoethanolamine
POPG: 1-palmitoyl-2-oleoyl-*sn*-glycero-3-phosphatidylglycerol
DPPG: 1,2-dipalmitoyl-*sn*-glycero-3-phosphatidylglycerol
SLPE: 1-stearoyl-2-linoleoyl-*sn*-glycero-3-phosphoethanolamine
HG: hourglass
IF: inward-open
OC: occluded
OF: outward-open

