## Supplementary File for "Impact of increased membrane realism on conformational sampling of proteins"

[†]Department of Chemical & Biomolecular Engineering, [‡]Department of Chemistry, [§]Center for Biophysics and Computational Biology, [‖]Beckman Institute for Advanced Science and Technology, [#]Cancer Center at Illinois, [⊥]Center for Digital Agriculture, [¶]Department of Plant Biology, [□]National Center for Supercomputing Applications, and [×]NIH Center for Macromolecular Modeling and Bioinformatics, University of Illinois at Urbana-Champaign, Urbana, IL 61801, United States

[a]Independent Researcher.

### Table of Contents

#### Figures and Tables

Table S1 Maximum Complexity membrane composition – 64 lipids S-4

Table S2 Maximum Complexity membrane composition – 128 lipids S-5

Table S3 Maximum Complexity membrane composition – 256 lipids S-6

Table S4 Maximum Complexity membrane composition – 512 lipids S-7

Table S5 Top10 Complexity membrane composition – 256 lipids S-8

Table S6 van der Waals Interaction energy analysis between annular lipid species and residues lining putative lipid binding site S-9

Figure S1 Top10 Complexity membrane composition – lipid structures and ratios S-10

Figure S2 SCD order parameters for carbon chain 1 of all phospholipids from Maximum Complexity simulations S-11

Figure S3 SCD order parameters for carbon chain 2 of all phospholipids from Maximum Complexity simulations S-12

Figure S4 SCD order parameters for carbon chain 1 of all phospholipids from Top10 Complexity simulations S-13

Figure S5 SCD order parameters for carbon chain 2 of all phospholipids from Top10 Complexity simulations S-14

Figure S6 Headgroup lateral tilt angle of all lipids shared by Maximum and Top10 Complexity membranes S-15

Figure S7 Headgroup axial tilt angle of all lipids shared by Maximum and Top10 Complexity membranes S-16

Figure S8 Sterol species electron density S-17

Figure S9 16:0/18:2 phospholipid species electron density S-18

Figure S10 18:2/18:2 phospholipid species electron density S-19

Figure S11 18:2/18:3 phospholipid species electron density S-20

Figure S12 16:0/18:1 phospholipid species electron density S-21

Figure S13 Water density S-22

Figure S14 MSM-weighted OsSWEET2b adaptive sampling error S-23

Figure S15 One-dimensional representation of OsSWEET2b gating dynamics S-24

Figure S16 Comparative landscape with previously published data of OsSWEET2b dynamics embedded in a POPC membrane S-25

Figure S17 OsSWEET2b annular shell composition graphs S-26

Figure S18 OsSWEET2b annular shell snapshots S-27

**Table S1.** Maximum Complexity membrane composition – 64 lipids

| | Upper Leaflet | Lower Leaflet |
|---|---|---|
| **Sterols** | 22 | 19 |
| *Beta-sitosterol (SITO)* | 15 | 16 |
| *Stigmasterol (STIG)* | 7 | 3 |
| | **Upper Leaflet** | **Lower Leaflet** |
| **Phospholipids** | 12 | 21 |
| *PC* | | |
| *16:0/18:2 (PLPC)* | 3 | 6 |
| *18:2/18:2 (DLiPC)* | 1 | 2 |
| *18:2/18:3 (LLPC)* | 1 | 2 |
| | **Upper Leaflet** | **Lower Leaflet** |
| *PE* | | |
| *16:0/18:2 (PLPE)* | 4 | 6 |
| *18:2/18:2 (DLiPE)* | 1 | 2 |
| *18:2/18:3 (LLPE)* | 1 | 1 |
| | **Upper Leaflet** | **Lower Leaflet** |
| *PG* | | |
| *16:0/18:2 (PLPG)* | 1 | 2 |

**Table S2.** Maximum Complexity membrane composition – 128 lipids

| | Upper Leaflet | Lower Leaflet |
|---|---|---|
| **Sterols** | 43 | 19 |
| *Beta-sitosterol (SITO)* | 29 | 13 |
| *Stigmasterol (STIG)* | 14 | 6 |
| | **Upper Leaflet** | **Lower Leaflet** |
| **Phospholipids** | 32 | 43 |
| *PC* | | |
| *16:0/18:1 (POPC)* | 1 | 1 |
| *16:0/18:2 (PLPC)* | 6 | 11 |
| *18:2/18:2 (DLiPC)* | 2 | 4 |
| *18:2/18:3 (LLPC)* | 2 | 4 |
| | **Upper Leaflet** | **Lower Leaflet** |
| *PE* | | |
| *16:0/18:2 (PLPE)* | 12 | 12 |
| *18:2/18:2 (DLiPE)* | 4 | 4 |
| *18:2/18:3 (LLPE)* | 3 | 3 |
| | **Upper Leaflet** | **Lower Leaflet** |
| *PG* | | |
| *16:0/18:2 (PLPG)* | 2 | 4 |

**Table S3.** Maximum Complexity membrane composition – 256 lipids

| | Upper Leaflet | Lower Leaflet |
|---|---|---|
| **Sterols** | 86 | 37 |
| *Beta-sitosterol (SITO)* | 58 | 25 |
| *Stigmasterol (STIG)* | 28 | 12 |
| | **Upper Leaflet** | **Lower Leaflet** |
| **Phospholipids** | 46 | 83 |
| *PC* | | |
| *16:0/18:1 (POPC)* | 2 | 2 |
| *16:0/18:2 (PLPC)* | 11 | 20 |
| *18:0/18:0 (DSPC)* | 1 | 2 |
| *18:1/18:1 (DOPC)* | 1 | 1 |
| *18:2/18:2 (DLiPC)* | 4 | 8 |
| *18:2/18:3 (LLPC)* | 3 | 8 |
| | **Upper Leaflet** | **Lower Leaflet** |
| *PE* | | |
| *16:0/18:1 (POPE)* | 1 | 0 |
| *16:0/18:2 (PLPE)* | 13 | 20 |
| *18:2/18:2 (DLiPE)* | 4 | 8 |
| *18:2/18:3 (LLPE)* | 2 | 6 |
| | **Upper Leaflet** | **Lower Leaflet** |
| *PG* | | |
| *16:0/16:0 (DPPG)* | 0 | 1 |
| *16:0/18:1 (POPG)* | 0 | 1 |
| *16:0/18:2 (PLPG)* | 4 | 6 |

**Table S4.** Maximum Complexity membrane composition – 512 lipids

| | Upper Leaflet | Lower Leaflet |
|---|---|---|
| **Sterols** | 173 | 74 |
| *Beta-sitosterol (SITO)* | 117 | 50 |
| *Stigmasterol (STIG)* | 56 | 24 |
| | **Upper Leaflet** | **Lower Leaflet** |
| **Phospholipids** | 93 | 172 |
| *PC* | | |
| *16:0/18:1 (POPC)* | 3 | 4 |
| *16:0/18:2 (PLPC)* | 22 | 40 |
| *18:0/18:0 (DSPC)* | 1 | 3 |
| *18:1/18:1 (DOPC)* | 2 | 1 |
| *18:2/18:2 (DLiPC)* | 9 | 17 |
| *18:2/18:3 (LLPC)* | 6 | 15 |
| | **Upper Leaflet** | **Lower Leaflet** |
| *PE* | | |
| *16:0/18:1 (POPE)* | 2 | 1 |
| *16:0/18:2 (PLPE)* | 24 | 47 |
| *18:0/18:2 (SLPE)* | 1 | 2 |
| *18:2/18:2 (DLiPE)* | 7 | 15 |
| *18:2/18:3 (LLPE)* | 4 | 11 |
| | **Upper Leaflet** | **Lower Leaflet** |
| *PG* | | |
| *16:0/16:0 (DPPG)* | 1 | 1 |
| *16:0/18:1 (POPG)* | 1 | 2 |
| *16:0/18:2 (PLPG)* | 10 | 13 |

**Table S5.** Top10 Complexity membrane composition – 256 lipids

| | Upper Leaflet | Lower Leaflet |
|---|---|---|
| **Sterols** | 75 | 47 |
| *Beta-sitosterol (SITO)* | 51 | 32 |
| *Stigmasterol (STIG)* | 24 | 15 |
| | **Upper Leaflet** | **Lower Leaflet** |
| **Phospholipids** | 57 | 77 |
| *PC* | | |
| *16:0/18:1 (POPC)* | 1 | 1 |
| *16:0/18:2 (PLPC)* | 15 | 20 |
| *18:2/18:2 (DLiPC)* | 6 | 8 |
| *18:2/18:3 (LLPC)* | 4 | 8 |
| | **Upper Leaflet** | **Lower Leaflet** |
| *PE* | | |
| *16:0/18:2 (PLPE)* | 17 | 24 |
| *18:2/18:2 (DLiPE)* | 5 | 7 |
| *18:2/18:3 (LLPE)* | 2 | 5 |
| | **Upper Leaflet** | **Lower Leaflet** |
| *PG* | | |
| *16:0/18:2 (PLPG)* | 7 | 4 |

**Table S6.** van der Waals Interaction energy analysis between annular lipid species and residues lining putative lipid binding site. Simulation states taken from Maximum and Top10 Complexity OsSWEET2b simulations within the intracellular versus extracellular gating distance landscapes (taken from regions with extracellular gating distance between 5 and 6 Å, intracellular gating distance between 13.25 and 13.75 Å). A “Full Binder” refers to an instance where a lipid species primarily occupies the OsSWEET2b putative binding site. A “Partial Binder” refers to an instance where a lipid species interacts with residues from the putative binding site but does not assume the same pose as a “Full Binder”.

| | | Residues found in putative lipid binding site | | | | | | | |
|---|---|---|---|---|---|---|---|---|---|
| | | Ile17 | Phe18 | Leu24 | Val27 | Thr28 | Tyr168 | Leu171 | Leu175 |
| ***Full Binders*** | | | | | | | | | |
| *Maximum* | PLPE (16:0/18:2) | -0.7055 ± 0.1241 | -1.9568 ± 0.1989 | -0.4967 ± 0.2129 | -0.0498 ± 0.0214 | -0.1560 ± 0.0485 | -5.9336 ± 0.4085 | -1.7764 ± 0.1595 | -1.082 ± 0.1256 |
| | DLIPE (18:2/18:2) | -1.4530 ± 0.1750 | -1.1058 ± 0.1444 | -3.5800 ± 0.2103 | -1.5919 ± 0.1643 | -2.5966 ± 0.2210 | -5.0433 ± 0.2923 | -0.5063 ± 0.0827 | -0.2691 ± 0.0505 |
| | LLPC (18:2/18:3) | -0.9160 ± 0.0528 | -1.0608 ± 0.0595 | -0.4810 ± 0.0626 | -0.2511 ± 0.0391 | -0.4738 ± 0.0573 | -2.6409 ± 0.1406 | -0.8298 ± 0.0515 | -0.5119 ± 0.0359 |
| | SITO | -- | -- | -- | -- | -- | -- | -- | -- |
| *Top10* | PLPC (16:0/18:2) | -1.1426 ± 0.1235 | -0.8783 ± 0.1062 | -4.079 ± 0.1742 | -2.5604 ± 0.1614 | -2.8729 ± 0.1981 | -4.6327 ± 0.2490 | -0.6691 ± 0.0997 | -0.2966 ± 0.0493 |
| | DLIPE (18:2/18:2) | -0.6487 ± 0.1642 | -2.2963 ± 0.2529 | -- | -- | -- | -4.1511 ± 0.3895 | -2.7956 ± 0.1743 | -2.0288 ± 0.1987 |
| | LLPC (18:2/18:3) | -- | -- | -0.0230 ± 0.0160 | -0.0425 ± 0.0254 | -0.0015 ± 0.0010 | -- | -- | -- |
| | SITO | -0.6115 ± 0.0644 | -1.5650 ± 0.0819 | -1.3331 ± 0.1303 | -0.8114 ± 0.0943 | -0.9045 ± 0.1090 | -4.1985 ± 0.1422 | -1.4661 ± 0.0927 | -0.9775 ± 0.0779 |
| ***Partial Binders*** | | | | | | | | | |
| *Maximum* | SITO | -- | -0.9156 ± 0.1672 | -2.2346 ± 0.0376 | -2.025 ± 0.0473 | -0.7367 ± 0.0228 | -- | -0.6545 ± 0.0749 | -0.7729 ± 0.0796 |
| *Top10* | SITO | -0.0614 ± 0.0062 | -0.7509 ± 0.0765 | -1.8331 ± 0.1512 | -2.6128 ± 0.0698 | -0.2891 ± 0.0268 | -1.1435 ± 0.0910 | -1.5459 ± 0.0818 | -0.9292 ± 0.0653 |

0 to -1 kcal/mol
-1 to -2 kcal/mol
-2 to -3 kcal/mol
-3 to -4 kcal/mol
-4 to -5 kcal/mol
-5 to -6 kcal/mol

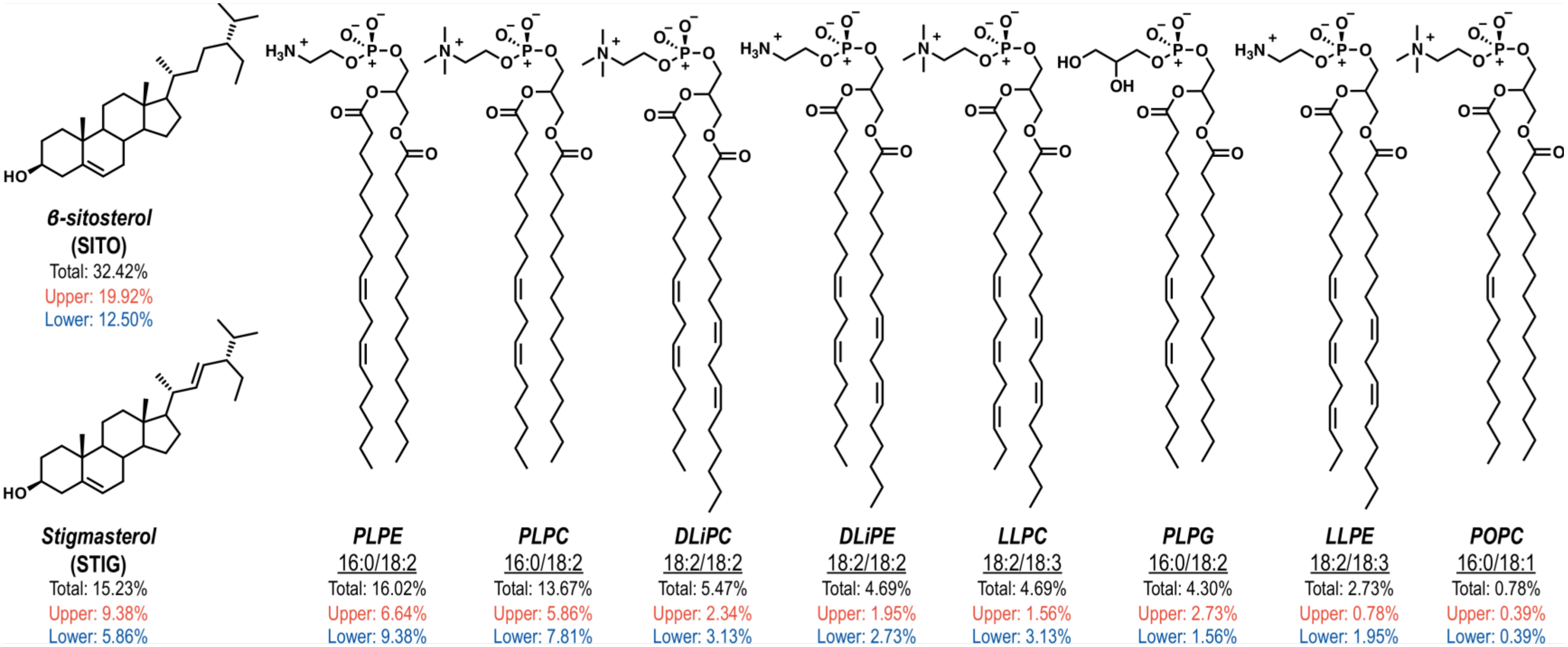


**Figure S1.** Top10 Complexity membrane composition – lipid structures and ratios

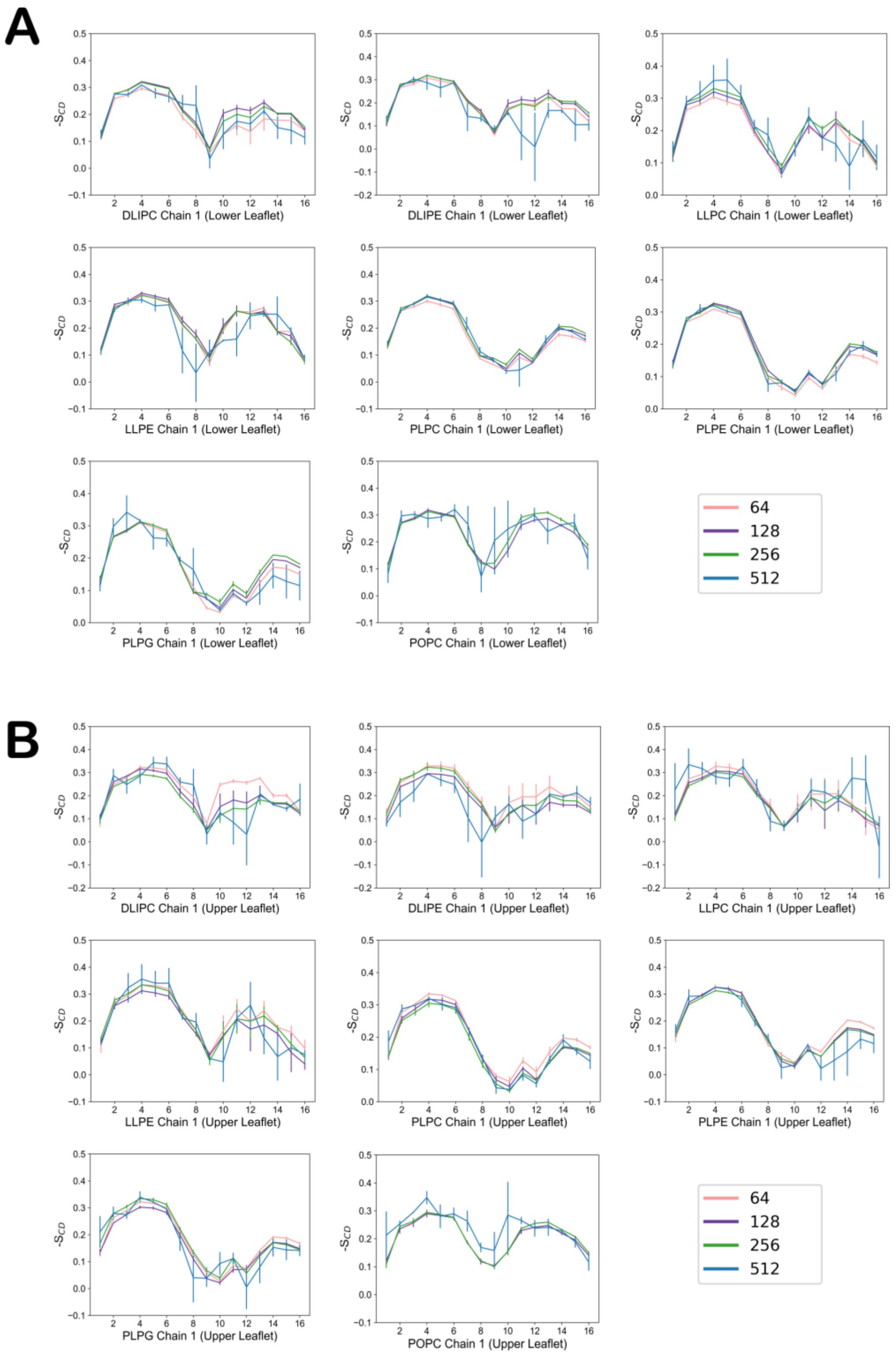


**Figure S2.** SCD order parameters for carbon chain 1 of all phospholipids from Maximum Complexity simulations. (A) Carbon chain 1, lower leaflet. (B) Carbon chain 1, upper leaflet.

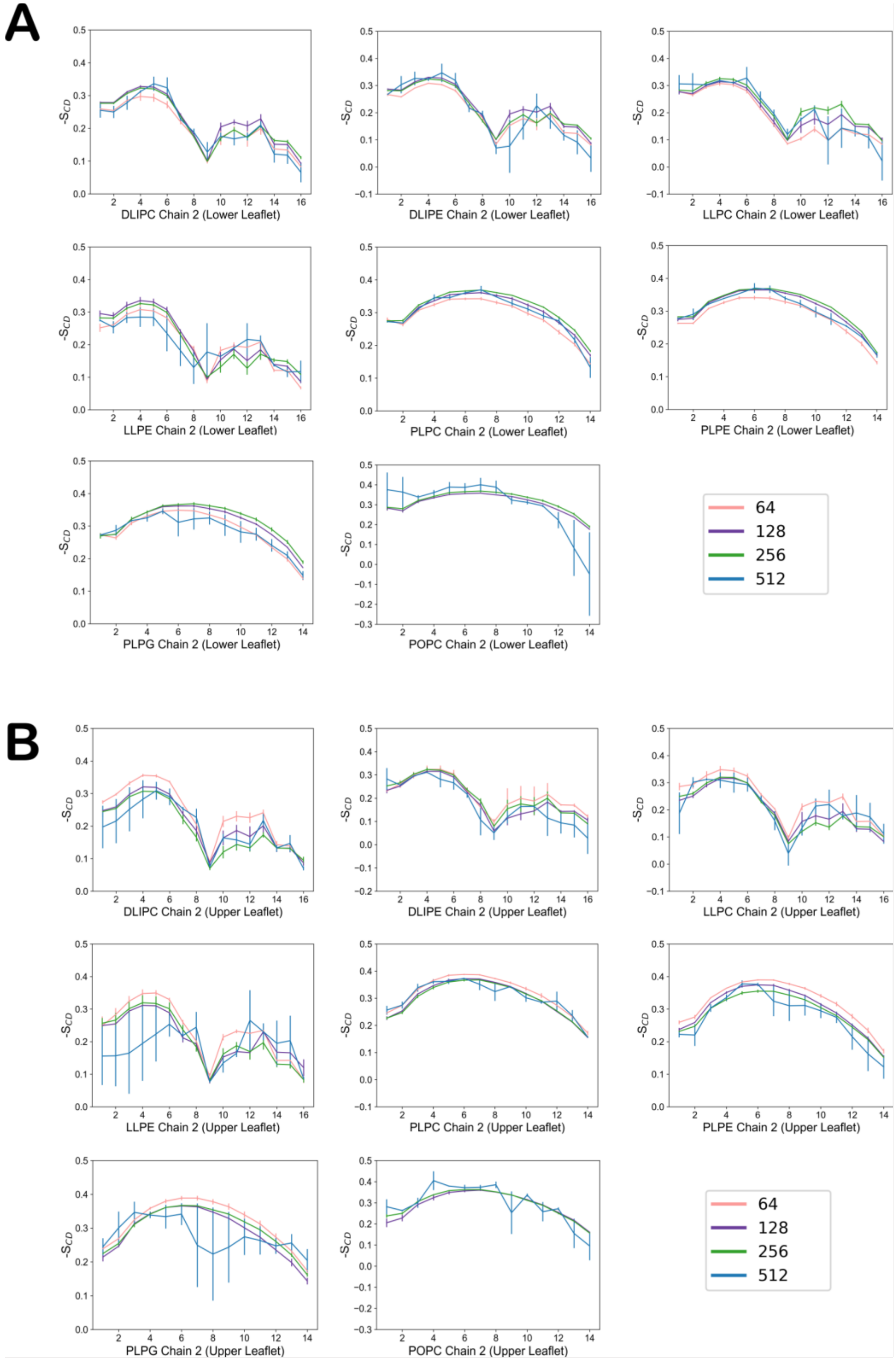


**Figure S3.** SCD order parameters for carbon chain 2 of all phospholipids from Maximum Complexity simulations. (A) Carbon chain 2, lower leaflet. (B) Carbon chain 2, upper leaflet.

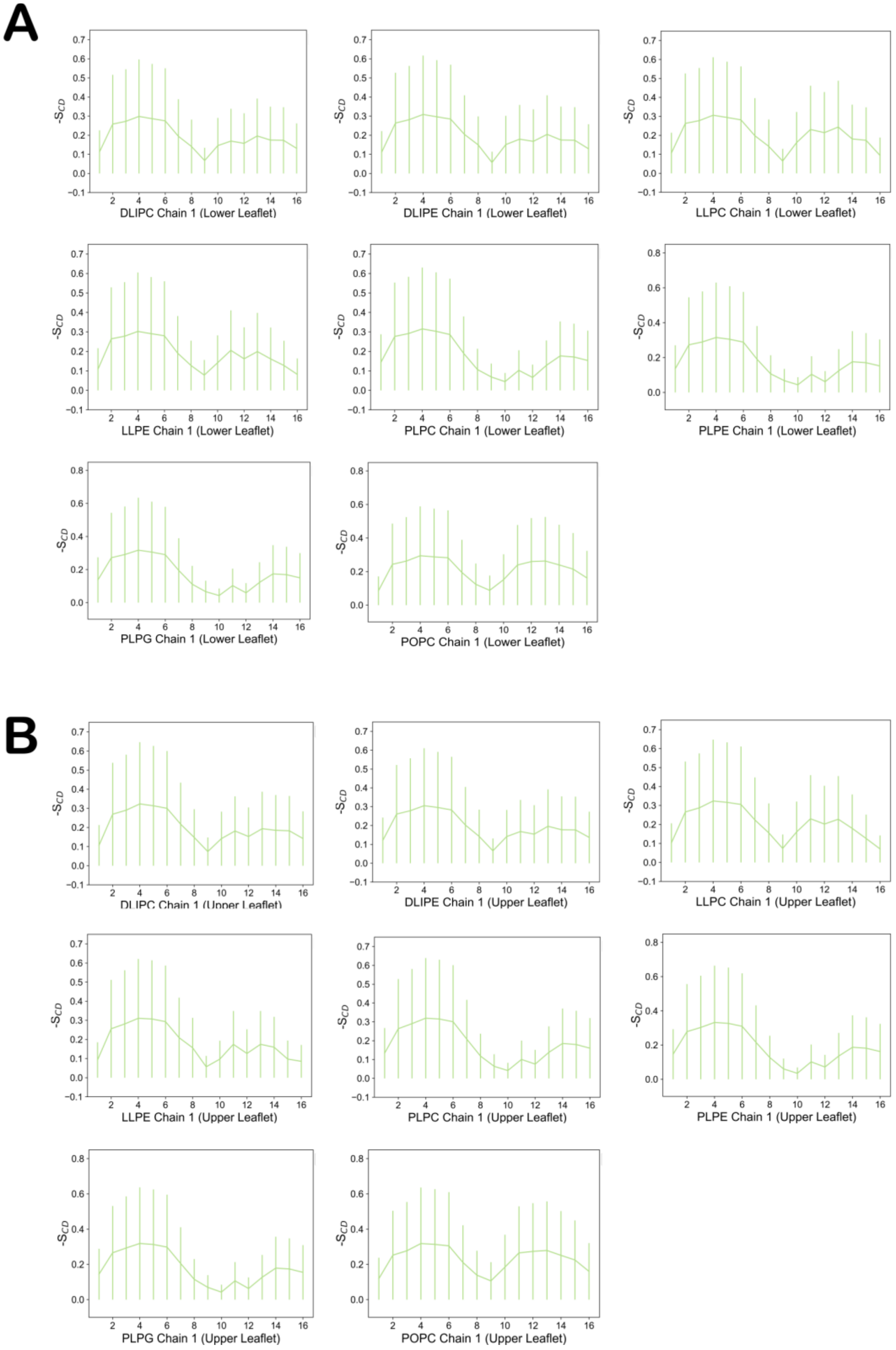


**Figure S4.** SCD order parameters for carbon chain 1 of all phospholipids from Top10 Complexity simulations. (A) Carbon chain 1, lower leaflet. (B) Carbon chain 1, upper leaflet.

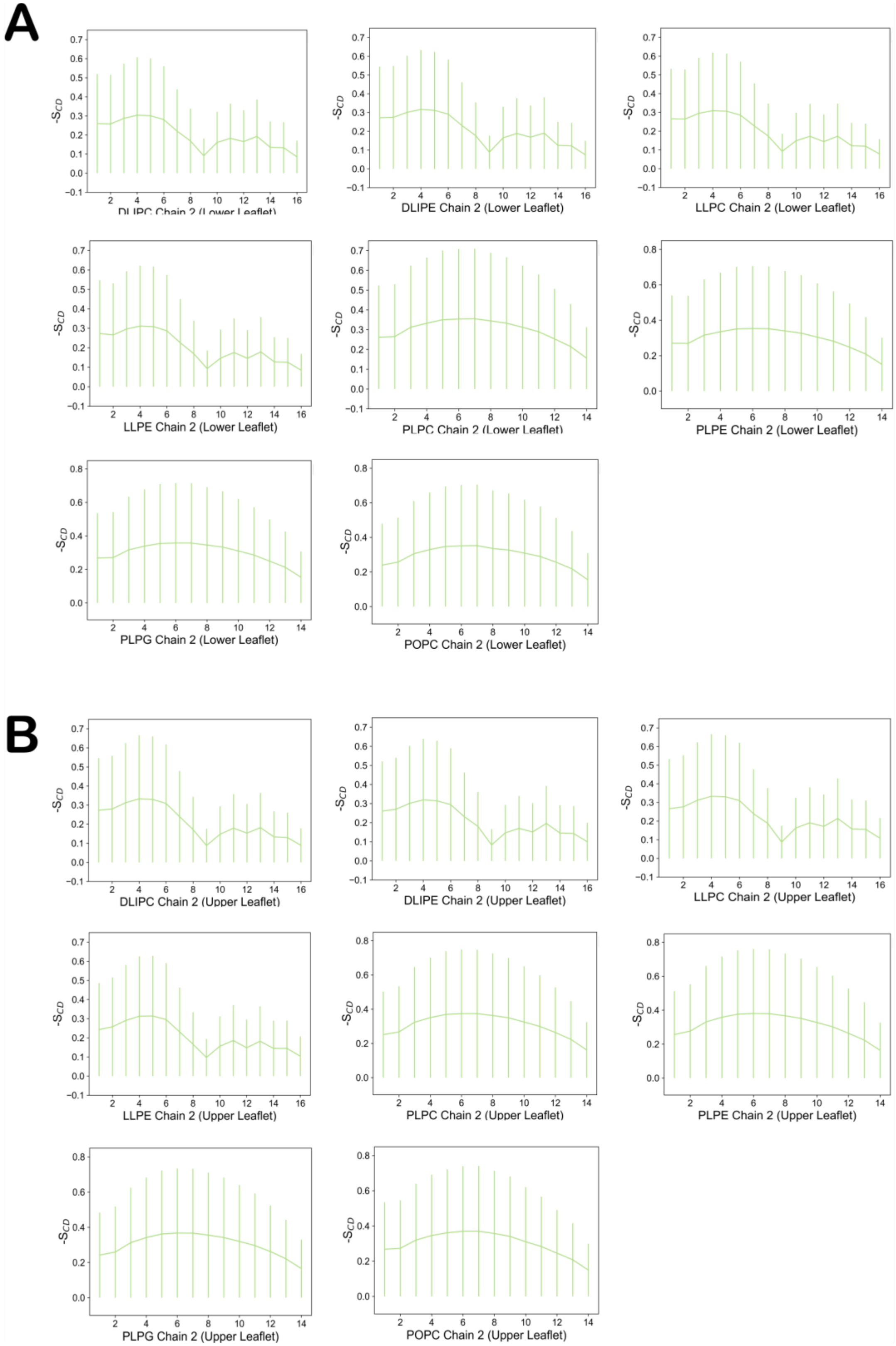


**Figure S5.** SCD order parameters for carbon chain 1 of all phospholipids from Top10 Complexity simulations. (A) Carbon chain 1, lower leaflet. (B) Carbon chain 1, upper leaflet.

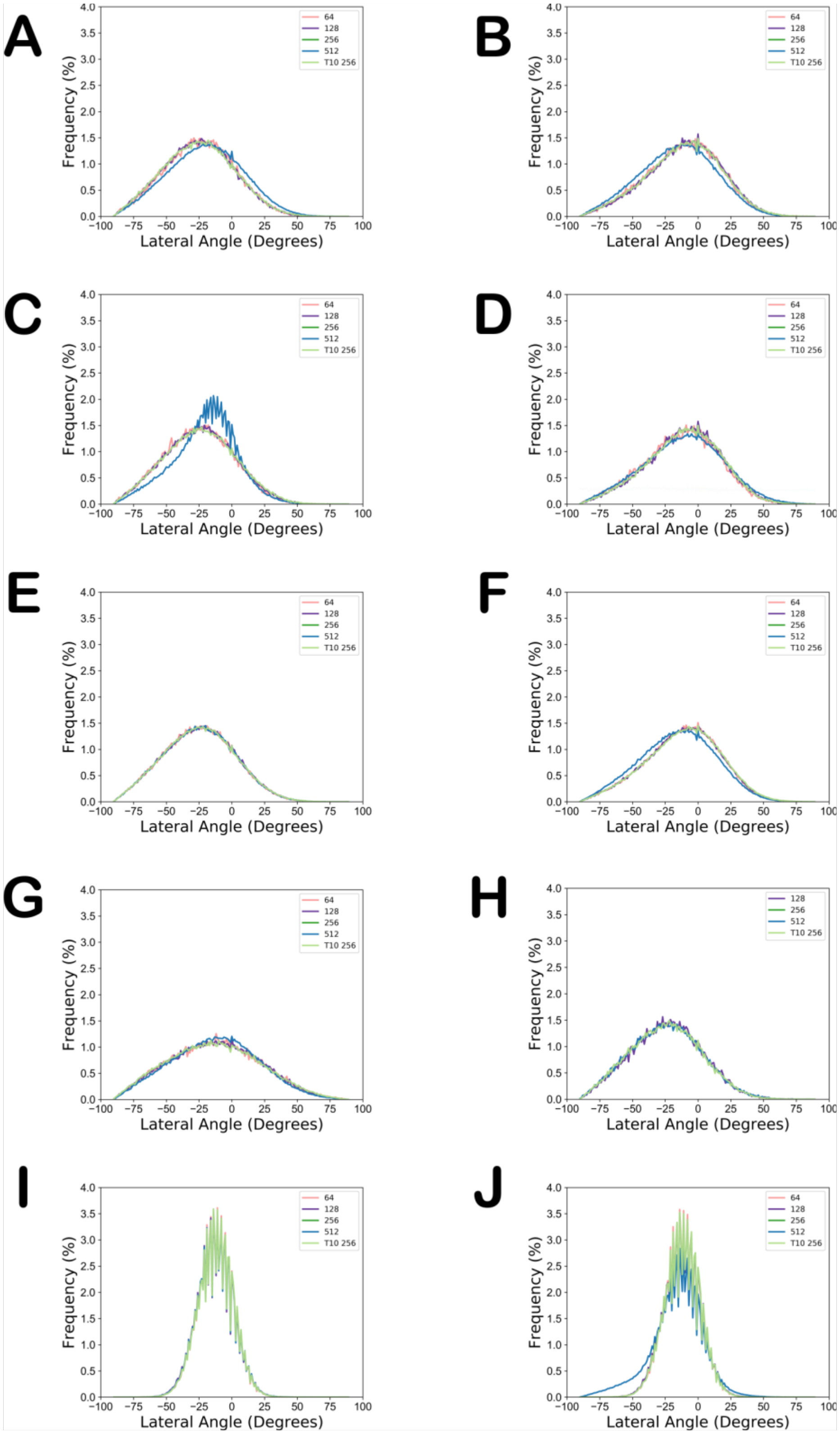


**Figure S6.** Headgroup lateral tilt angle of all lipids shared by Maximum and Top10 Complexity simulations. (A) DLiPC. (B) DLiPE. (C) LLPC. (D) LLPE. (E) PLPC. (F) PLPE. (G) PLPG. (H) POPC (I) SITO. (J) STIG.

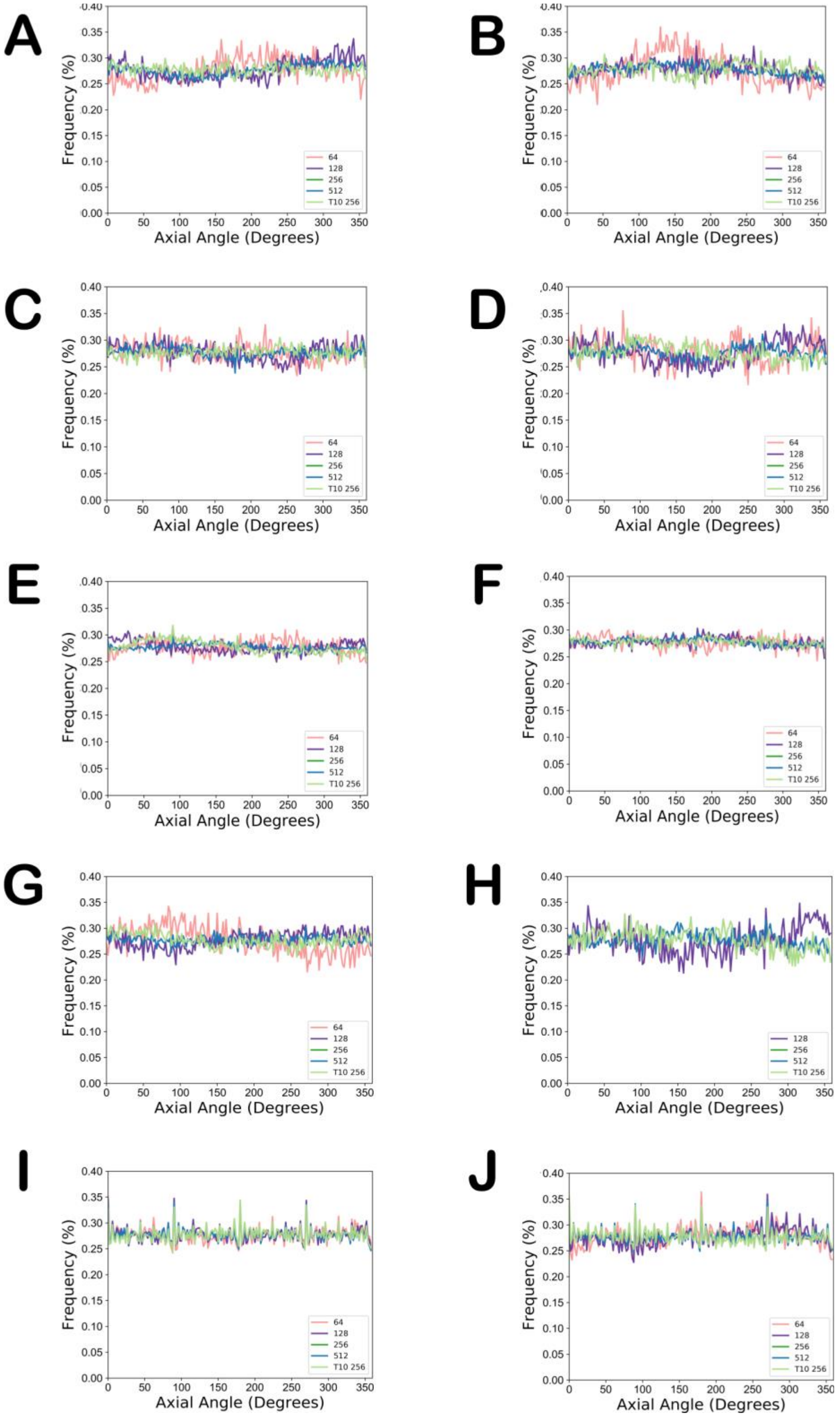


**Figure S7.** Headgroup axial tilt angle of all lipids shared by Maximum and Top10 Complexity simulations. (A) DLiPC. (B) DLiPE. (C) LLPC. (D) LLPE. (E) PLPC. (F) PLPE. (G) PLPG. (H) POPC (I) SITO. (J) STIG.

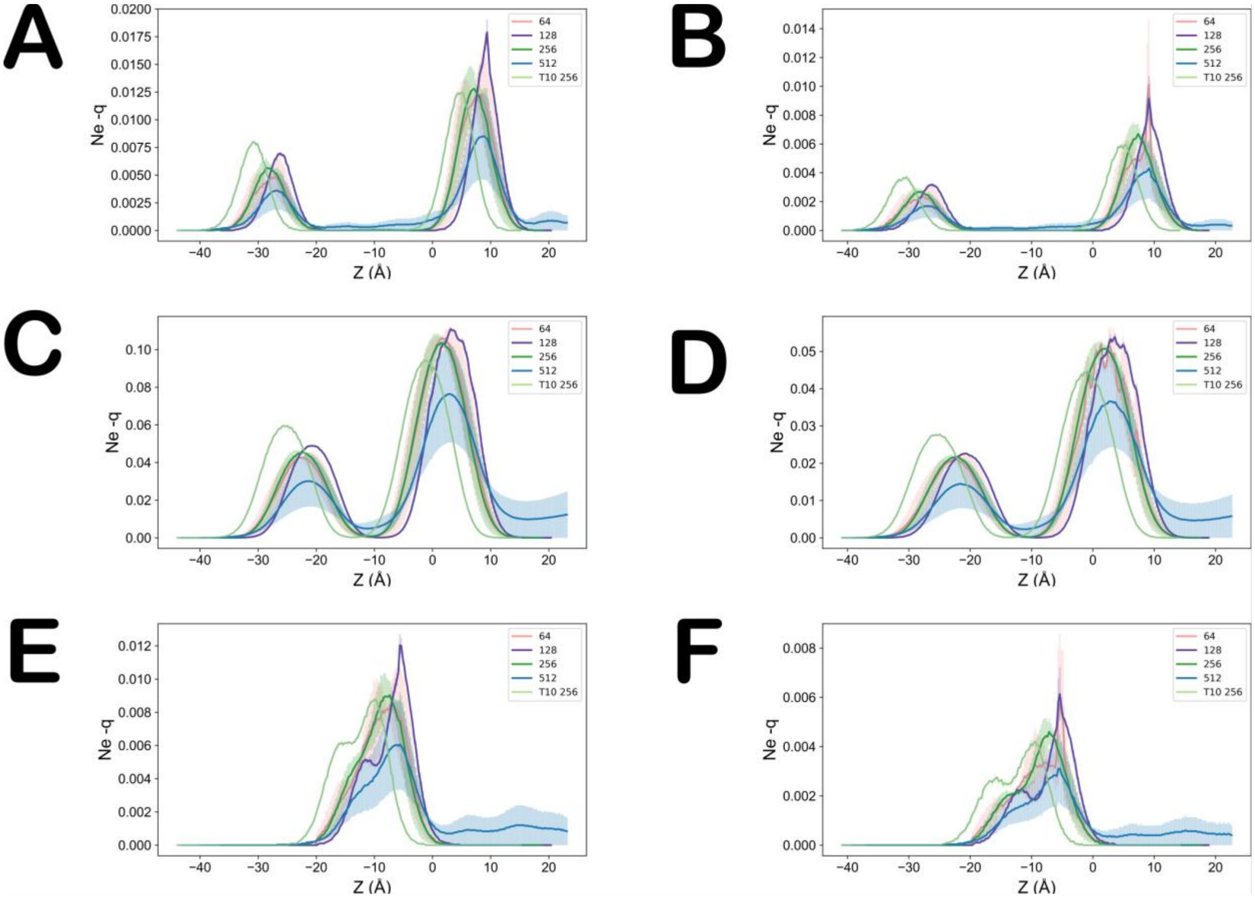


**Figure S8.** Sterol species electron density. (A) SITO O3 density. (B) STIG O3 density. (C) SITO rings density. (D) STIG rings density. (E) SITO C24 density. (F) STIG C24 density.

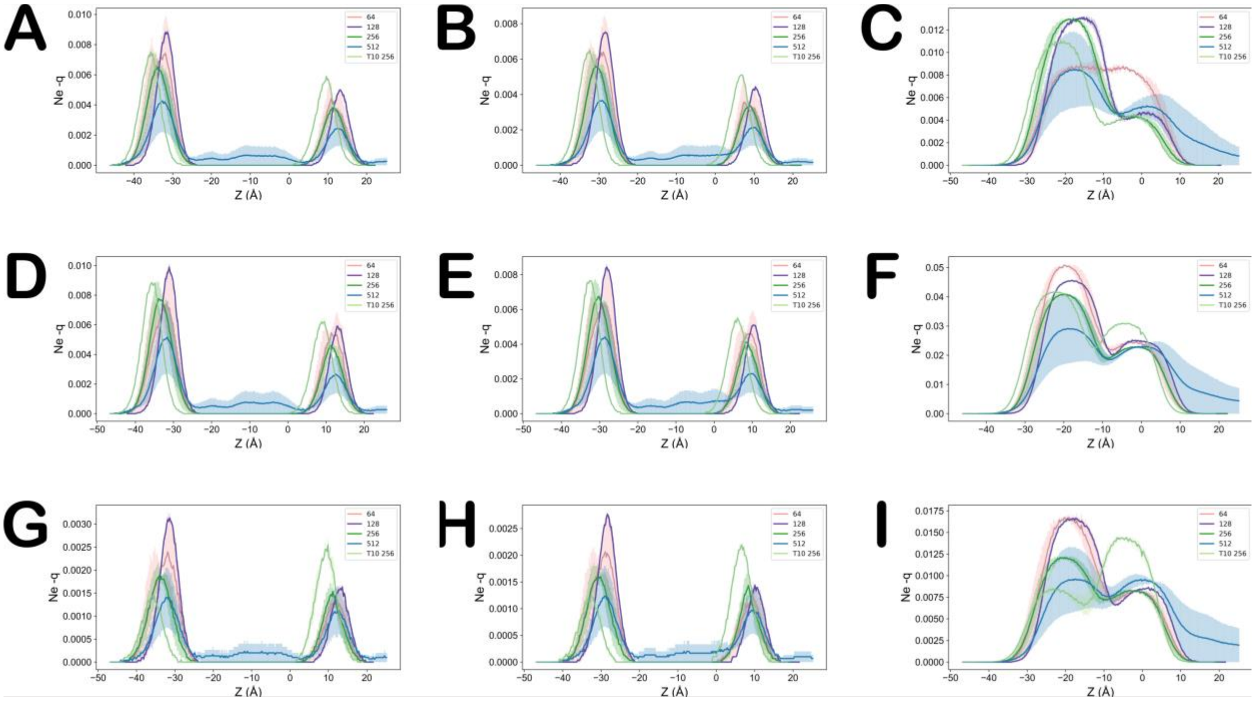


**Figure S9.** 16:0/18:2 phospholipid species electron density. (A) PLPC P density. (B) PLPC glycerol density. (C) PLPC tail density. (D) PLPE P density. (E) PLPE glycerol density. (F) PLPE tail density. (G) PLPG P density. (H) PLPG glycerol density. (I) PLPG tail density.

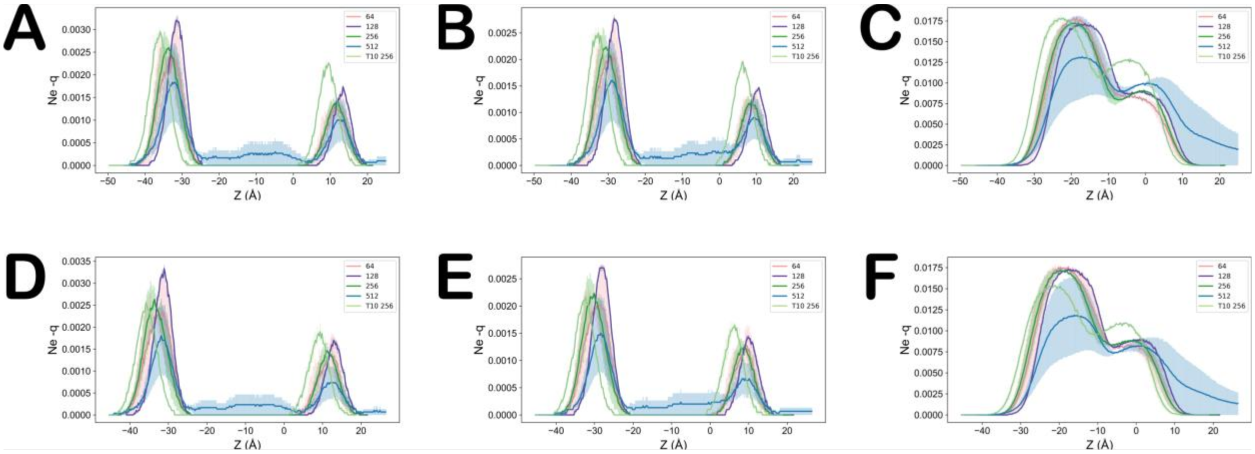


**Figure S10.** 18:2/18:2 phospholipid species electron density. (A) DLiPC P density. (B) DLiPC glycerol density. (C) DLiPC tail density. (D) DLiPE P density. (E) DLiPE glycerol density. (F) DLiPE tail density.

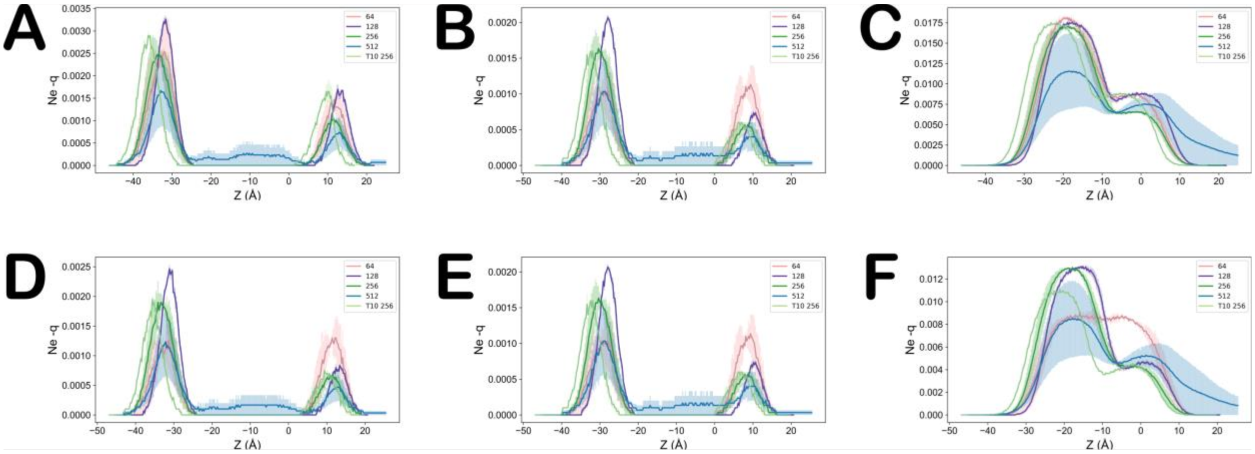

**Figure S11.** 18:2/18:3 phospholipid species electron density. (A) LLPC P density. (B) LLPC glycerol density. (C) LLPC tail density. (D) LLPE P density. (E) LLPE glycerol density. (F) LLPE tail density.

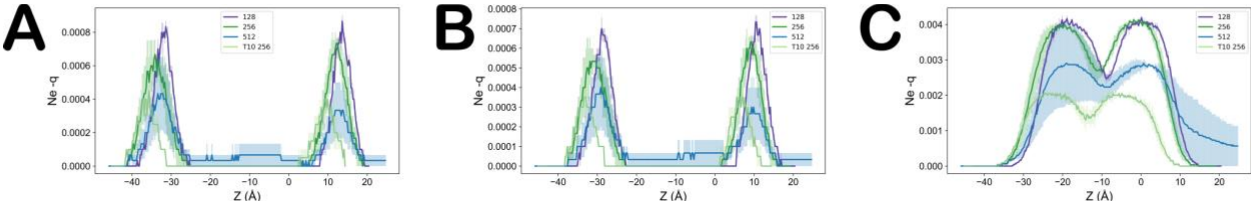


**Figure S12.** 16:0/18:1 phospholipid species electron density. (A) POPC P density. (B) POPC glycerol density. (C) POPC tail density.

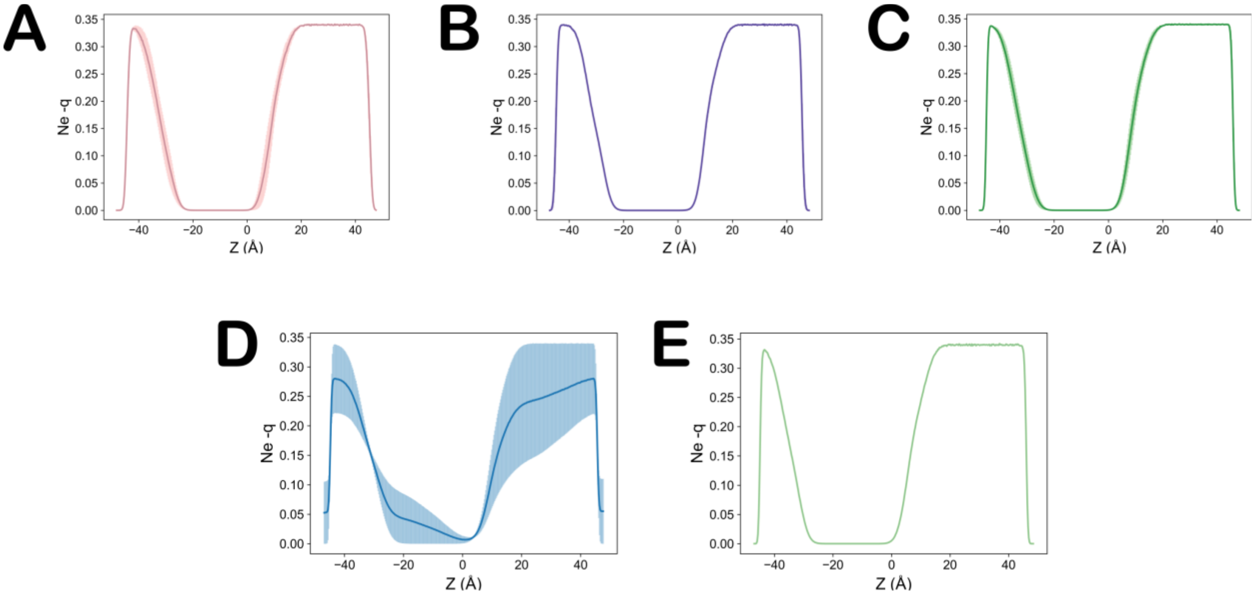


**Figure S13.** Water density. (A) WAT 64 from Maximum Complexity simulations. (B) WAT 128 from Maximum Complexity simulations. (C) WAT 256 from Maximum Complexity simulations. (D) WAT 512 from Maximum Complexity simulations. (E) WAT 256 from Top10 Complexity simulations.

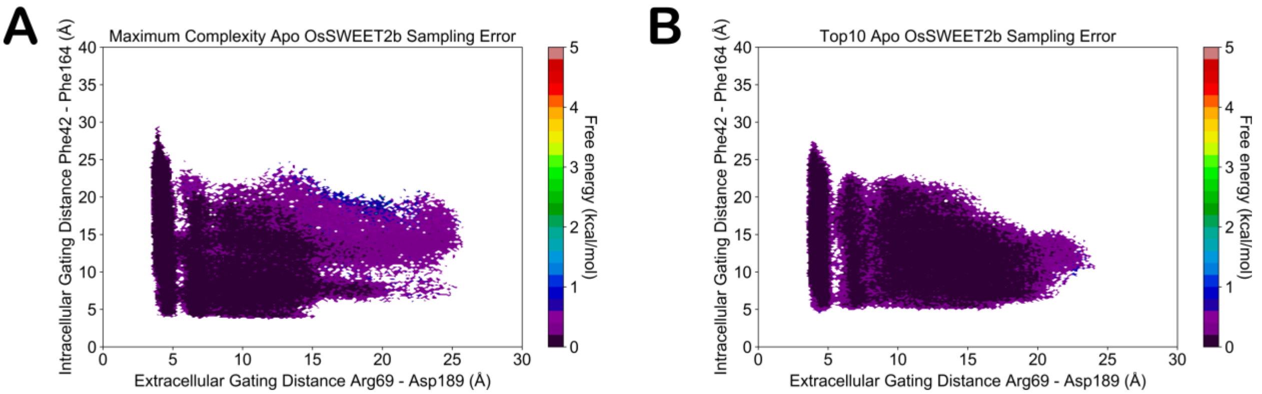


**Figure S14.** MSM-weighted OsSWEET2b adaptive sampling error. (A) Maximum Complexity OsSWEET2b gating dynamics error projection. (B) Top10 Complexity OsSWEET2b gating dynamics error projection.

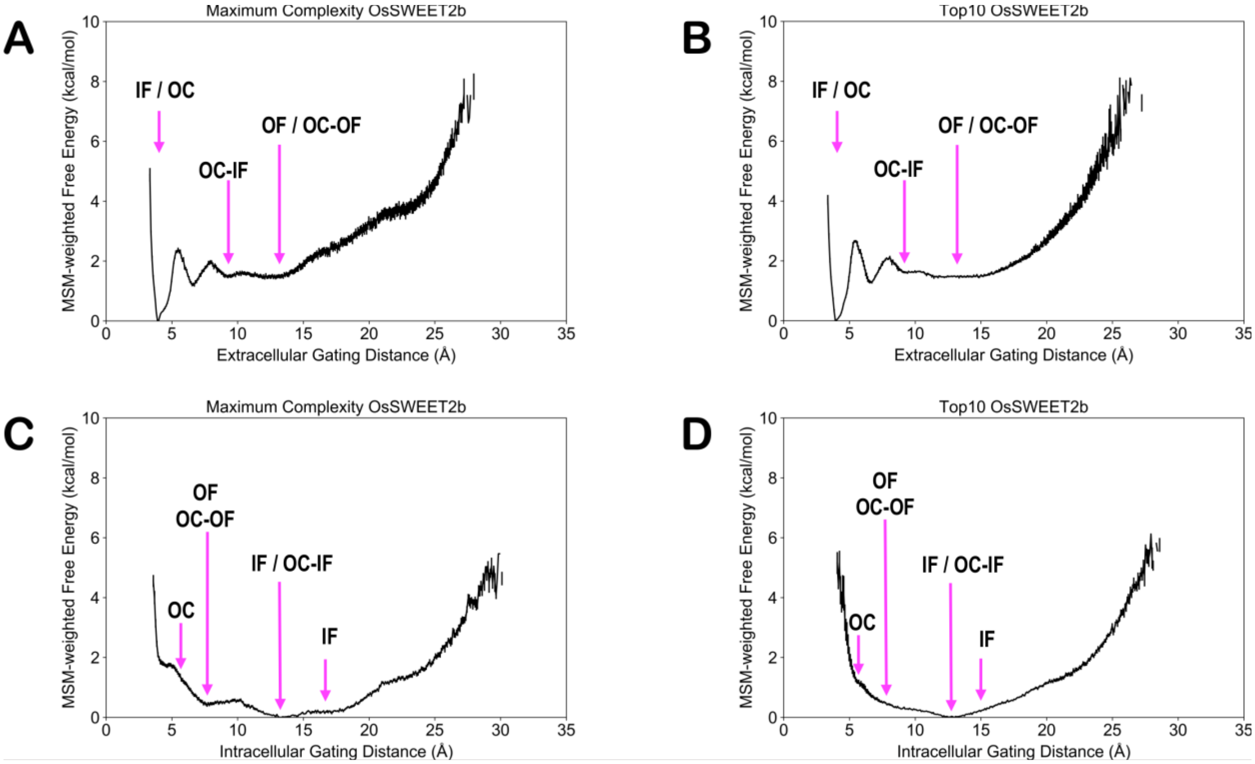


**Figure S15.** One-dimensional representation of OsSWEET2b gating dynamics. (A) Extracellular gating dynamics for Maximum Complexity OsSWEET2b construct. (B) Extracellular gating dynamics for Top10 Complexity OsSWEET2b construct. (C) Intracellular gating dynamics for Maximum Complexity OsSWEET2b construct. (D) Intracellular gating dynamics for Top10 Complexity OsSWEET2b construct.

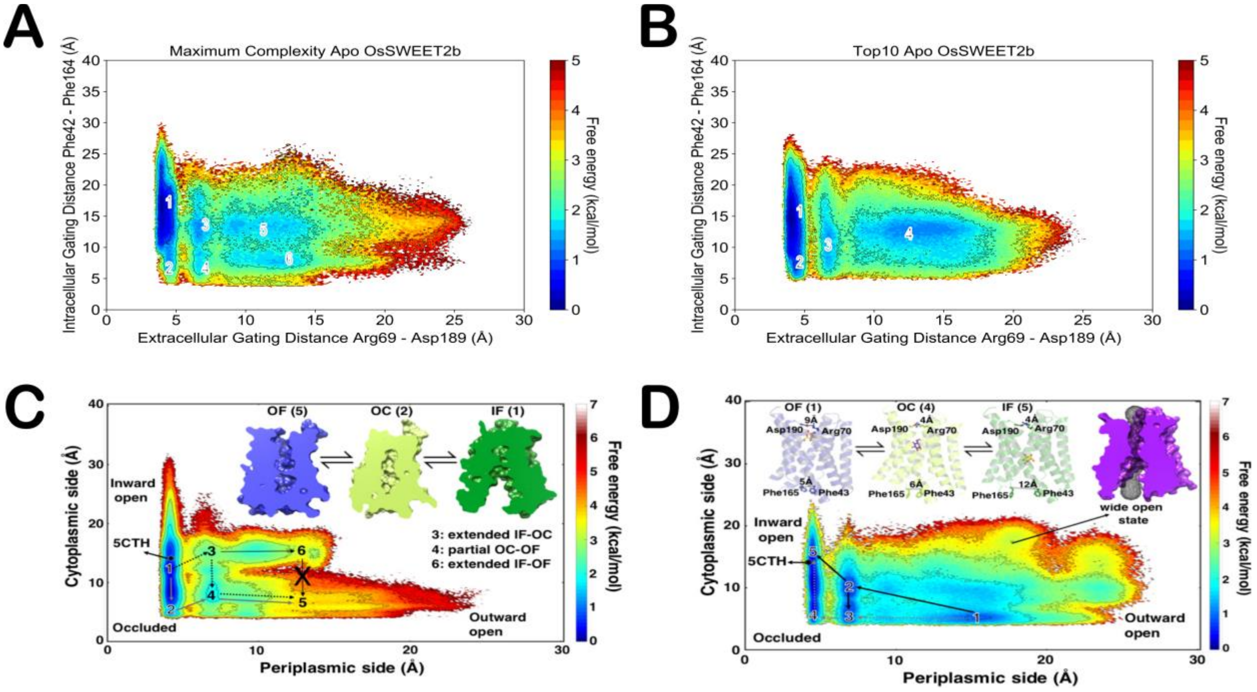


**Figure S16.** Comparative landscape with previously published data of OsSWEET2b dynamics embedded in a POPC membrane (main text citation Selvam, B.; Yu, Y-C.; Chen, L-Q.; Shukla, D. *ACS Cent. Sci.,* **2019**, *5*, 1085, https://pubs.acs.org/doi/full/10.1021/acscentsci.9b00252). Main text figures are colored with a different colormap to allow for more direct comparisons with prior data. (A) apo OsSWEET2b simulations done in the 256 Maximum Complexity membrane construct. (B) apo OsSWEET2b simulations done in the 256 Top10 Complexity membrane construct. (C) apo OsSWEET2b simulations done in a pure POPC bilayer. (D) holo OsSWEET2b simulations done in a pure POPC bilayer. Readers seeking further permissions to the material excerpted should contact ACS Publications Support.

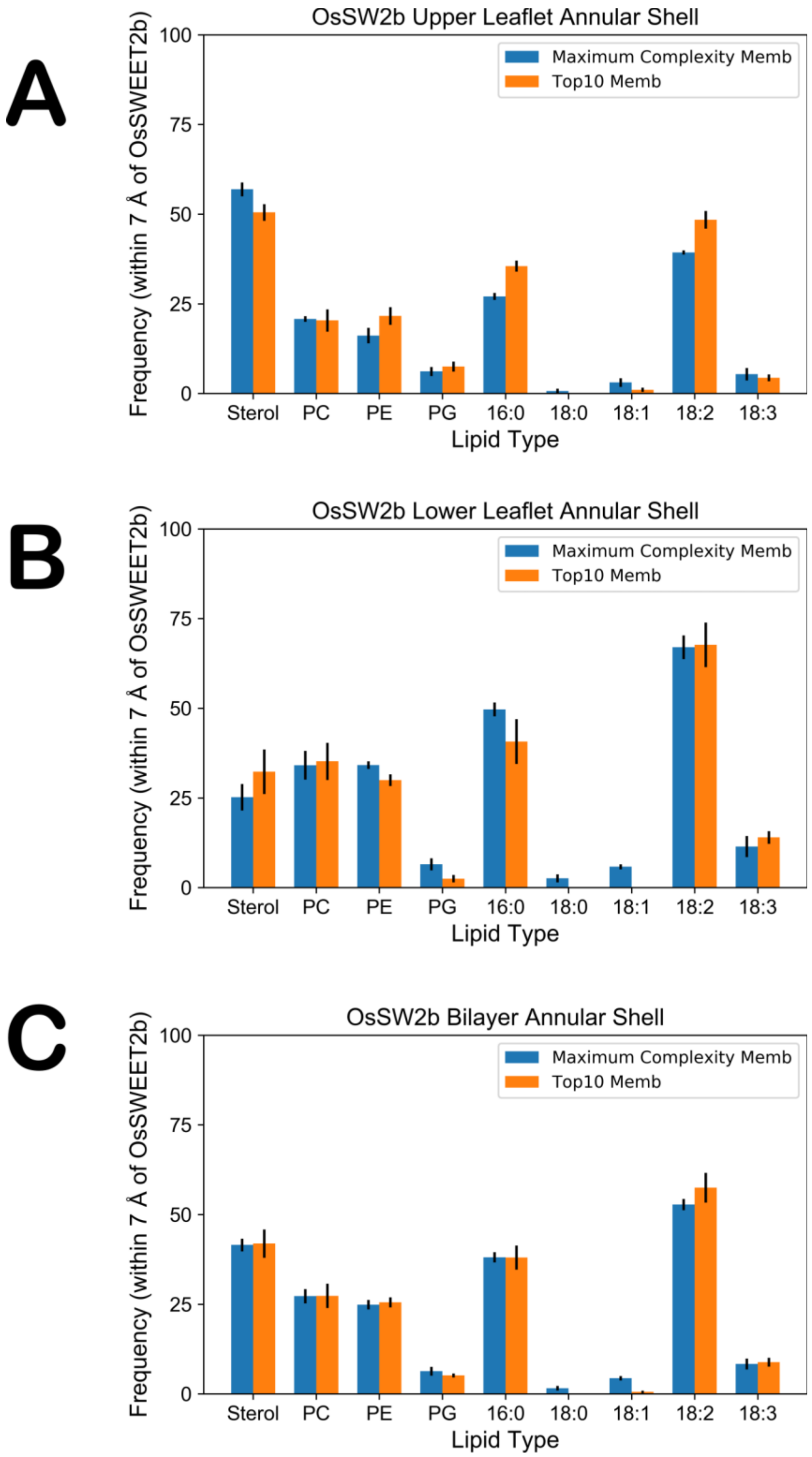


**Figure S17.** OsSWEET2b annular shell composition. Annular shell compositions are presented for the (A) upper and (B) lower leaflets, as well as for (C) the entire bilayer.

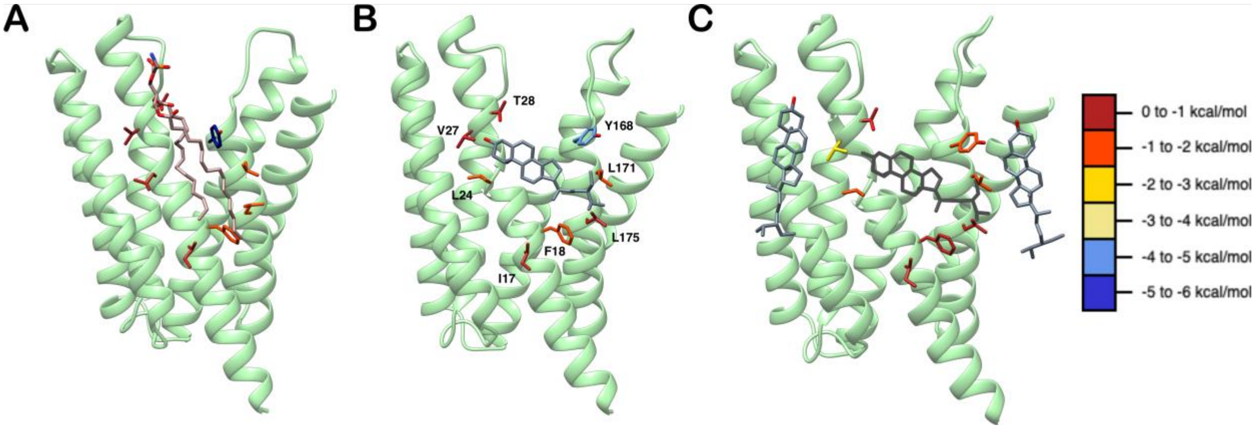


**Figure S18.** OsSWEET2b annular shell snapshots. Lipids bound to the putative binding lipid binding site are shown with residues colored with respect to their interaction energy values found in Supplemental Table 6. (A) PLPE bound in the "Full Binder" pose in OsSWEET2b Scale simulations. (B) SITO bound in the "Full Binder" pose in OsSWEET2b Top10 simulations. (C) SITO molecules bound in "Partial Binder" poses in OsSWEET2b Top10 simulations, with the "Full Binding" SITO greyed out. Here, the tendency of SITO molecules to aggregate around this site in Top10 simulations results in less stable interaction energies with putative binding site residues.
